# At the core of salinity: convergent and divergent transcriptome response pathways to neutral and alkaline salinity in natural populations of *Arabidopsis thaliana*

**DOI:** 10.1101/2023.02.05.527167

**Authors:** Maria Almira-Casellas, Sílvia Busoms, Laura Pérez-Martín, Glòria Escolà, Álvaro López-Valiñas, Antoni Garcia-Molina, Mercè Llugany, Charlotte Poschenrieder

## Abstract

More than 70% of land’s cultivated area is affected by alkaline salinity stress. As 98% of plants are glycophytes – unable to successfully reproduce under salinity – our previous research focused on comparative studies of *Arabidopsis thaliana* demes with differential performance under neutral and alkaline salinity *(neuSAL* and *alkSAL*) due to local adaptation processes. Here, an integrated analysis on leaf tissue was performed, including physiological indicators, nutritional status, endogenous phytohormonal concentration and transcriptome profiling, to further understand differences in molecular mechanisms underlying *neuSAL* and *alkSAL* responses. The results support that *alkSAL* is more detrimental to plant performance than *neuSAL* and indicate higher sensitivity to *alkSAL* in demes locally adapted to coastal siliceous soils. A decreased internal Fe use efficiency in coastal demes under *alkSAL* is proposed to be the driver of their enhanced sensitivity, and sequence variation at β-CA1 and α-CA1 locus is hypothesized to contribute to the imbalance of Fe homeostasis. Dissection on the down-regulated transcripts shared by *neuSAL* and *alkSAL* confirmed enhanced inhibition of central features on primary and secondary metabolism in coastal individuals under *alkSAL*. The cell wall and vacuolar β-galactosidase *BGAL4* was revealed as a candidate for conferring tolerance to *neuSAL* by favoring stress-regulated cell wall rearrangement, but not to *alkSAL*, probably due to pH-restricted enzymatic activity. In addition, differential modulation of endogenous phytohormonal cues was reported among salinity types and demes, by which higher alteration of the auxinic, ethylene and jasmonic acid signaling pathways was exerted by alkSAL but sustained ABA biosynthesis was detected only in coastal plants under neuSAL. Weighted correlation network analysis (WGCNA) confirmed the involvement of the identified candidate genes in co-expression modules significantly correlating with favorable responses to *neuSAL* and *alkSAL*. Overall, the present study provides useful insights into key targets for breeding improvement in alkaline saline soils.

## Introduction

Saline stress is a major agricultural constraint and its co-occurrence with alkalinity is an arising environmental concern, specially under arid and semiarid climates. Based on the soil map of the world by FAO/UNESCO, more than one million hectares of the Earth’s land are affected by alkaline salinity stress. This is the case of the Mediterranean region, where excess of soluble Na^+^, Ca^2+^, Mg^2+^, K^+^, CO_3_^2−^ and HCO^3-^ ions get to the superficial soil layers due to high evapotranspiration rates (Singh 2021). Besides the induction of osmotic, ion and ROS stress by neutral salinity, alkaline salinity exacerbates nutrient deficiency (Liu et al., 2021) and is reported to exert greater inhibition on plant germination, growth, photosynthesis, and root system activity (Guo et al., 2010; Marconi et al. 2013; Guo et al., 2017). Although there is increasing evidence that alkaline salinity is more detrimental for overall plant performance than neutral salinity, the mechanisms driving such synergistic effects are not yet clearly established. In this regard, more attention should be paid to the environmental factors that generate local-scale diversity patterns in the model plant *Arabidopsis thaliana*, since understanding natural genetic and phenotypic variation is essential for ecological and evolutionary studies (Bomblies et al., 2010).

Genetic differentiation occurs at all geographic scales in *A. thaliana*, and the study of local populations provides a valuable genetic tool for detecting adaptive traits (Bakker et al., 2006). To this end, differences in tolerance to salinity and moderate levels of carbonate as individual stress factors have been described among natural populations of *A. thaliana* in the NE region of Catalonia, Spain (Busoms et al., 2015; Terés et al., 2019). Although T6^CS^ and Ro2^CS^ were shown to perform better under salinity on siliceous soils than V1^PC^ and LM2^PC^, it was proven that tolerance to neutral salinity does not confer tolerance to alkaline salinity in the studied individuals, and that populations from soils with intermediate salinity and carbonate levels performed better under alkaline salinity stress due to enhanced phenotypic plasticity (Pérez-Martín et al., 2022). However, the transcriptomic basis of such differential response were not explored. Plants are exposed to the co-occurrence of biotic and abiotic stresses in their natural habitats but the effects of combined stresses at the genetic level are much less extensively reviewed than plant responses to single stress factors. Still, there are several studies targeting the existing interconnection of signaling pathways that regulate the coordination of plant responses to different stresses (Sharma et al., 2018; Hosseini et al., 2021; Georgii et al., 2017; Suzuki et al., 2016; Forieri et al., 2017; Rasmussen et al., 2013), whose results highlight that combinatorial stress triggers responses not evoked by single stressors. Despite the scarcity of studies on the effects of co-occurrence of salinization and alkalization at the transcriptomic level in *Arabidopsis*, transcriptome responses to alkaline salinity have been assessed in several crops. In cotton (*Gossypium hirsutum* L.), shared responses in transcripts involved in ion homeostasis were observed under NaCl and Na_2_CO_3_ conditions, while differential expression of genes involved in starch and sucrose metabolism was reported to be exclusive of Na_2_CO_3_ stress due to high pH (Zhang et al., 2018). Transcriptomics on alkalinity-tolerant and sensitive wheat varieties treated for nine days with 100 mM mixed salts (9:1 NaHCO_3_:Na_2_CO_3_ at pH 8.90) revealed enhanced ability of nutrient uptake, intracellular pH regulation and ROS scavenging activation as the main traits conferring tolerance to alkaline salinity (Sun et al., 2020). Similarly, recent transcriptome analysis in *Brassica napus* revealed higher expression of genes from carbohydrate metabolism, photosynthetic processes, ROS regulating, and response to salt stress attributing tolerance to alkaline salinity (Meng et al., 2017). Enhanced antioxidant capacity was also reported to be a major player in the tolerance response of alfalfa to saline-alkaline stress (An et al., 2016). In rice, alkaline tolerant and sensitive cultivars were compared at the transcriptomic level under 0.5% Na_2_CO_3_ solution and pH 11, and differentially expressed genes from plant hormone signal transduction pathway were primarily related to the tolerance response (Li et al., 2018).

Linking changes in gene expression profiles to functional consequences is the ultimate goal of transcriptomic analysis on stress tolerance-related traits (Yan et al., 2020). The combination of physiological studies with transcriptomic analyses provides a deeper understanding of biological processes (Yang et al., 2022) and, based on the available studies, some of which are reported above, nutrient homeostasis and changes in endogenous phytohormone levels are considered among the most important mechanisms conferring tolerance in the context of neutral and alkaline salinity. While maintaining nutrient homeostasis starts from ion sensing, uptake, transport, and activation of response mechanisms through regulation of gene networks (Farooq et al., 2018), the alteration in the content of phytohormone implies a specific crosstalk between endogenous plant growth regulators. This leads to the triggering of signal transduction pathways in response to external conditions and can affect germination (Atici et al., 2005), morphology of root and aerial structures (Mao et al., 2018, Wen et al., 2018, Song et al., 2019), and reproduction, specially flowering (Guo et al., 2019, Shu et al., 2018, Bao et al., 2019). These considerations reinforce the idea that differences in mineral nutrition and phytohormone profile orchestrate adaptive processes in populations grown on specific local environments by modulating their growth and reproductive responses.

The present study aims to identify the levels and functions of plant stress regulatory networks underlying two contrasted types of soil salinity: neutral and alkaline. To this end, we seeked for shared and exclusive *Arabidopsis* responses to both stress conditions among genotypes locally adapted to either siliceous (neutral) or calcareous (alkaline) saline substrates.. Four *A. thaliana* local populations (demes) were subjected to neutral (*neuSAL*) and alkaline salinity (*alkSAL*) under the same light and temperature conditions. The physiological and transcriptomic responses of the 4 demes under each type of salinity were analyzed, and convergent and singular metabolic pathways involved in each type of salinity are described. Finally, co-expression network analysis correlating transcriptomics with physiological data were performed to identify hub pathways in shared and exclusive plant responses to neutral and alkaline salinity.

## Results & Discussion

### 1. Physiological responses of A. thaliana demes under neutral and alkaline salinity

To see if the adaptation to specific soils confer tolerance to neutral and alkaline salinity, two *A. thaliana* demes from coastal siliceous soils (cs), NaCl-rich and pH between 5.7-6.2 (T6^CS^ and Ro2^CS^); and 2 demes from pre-litoral calcareous soils (pa), considered non-saline, and with moderate carbonate content (10-15%) (V1^PC^, LM2^PC^) were selected. Plants were grown on potting mix soil irrigated with *neuSAL* (100 mM NaCl, pH 5.9) or *alkSAL* (85 mM NaCl; 15 mM NaHCO_3_, pH 8.3) for 2 weeks. The physiological, nutritional, and phytohormonal responses of the studied demes were assessed and multivariate analyses of all measured variables were performed to establish the association between traits and plant stress tolerance. Figures S1 and S2 detail the geographic location and source of the studied demes and the study workflow, respectively.

#### Demes from coastal siliceous soils display reduced growth and photosynthesis under alkaline salinity

Overall, *neuSAL* and *alkSAL* caused 50% decrease in relative rosette diameter (RD) [(RD_Treatment_)/ (RD_Control_)] regardless of deme and treatment, although LM2^PC^ showed significantly better growth maintenance than the other demes under *alkSAL* conditions (*p*-value ≤ 0.05, Tukey’s HSD) (Figure 1A; Dataset S1). The chlorophyll (Chl) fluorescence parameter, Fv/Fm, reflects the maximum quantum efficiency of PSII photochemistry in the dark-adapted state and has been widely used for the detection of early stress responses in plants. Under control and *neuSAL* conditions, no significant differences in Fv/Fm among demes were observed and mean Fv/Fm values were comprised in the reading considered as optimal (between 0.79 and 0.84) for stress free plants (Maxwell and Johnson 2004). Under *alkSAL*, T6^CS^ displayed a dramatic decrease in F_v_/F_m_ values (mean F_v_/F_m_ = 0.520) when compared to the rest, while V1^PC^ (mean F_v_/F_m_ = 0.770) and LM2^PC^ (mean F_v_/F_m_ = 0.820) showed better F_v_/F_m_ maintenance (*p*-value ≤ 0.05, Tukey’s HSD). Contrarily, under *neuSAL*, T6^CS^ and Ro2^CS^ (0.860 and 0.810, respectively) displayed significantly higher F_v_/F_m_ values than V1^PC^ (0.740) and LM2^PC^ (0.730) (Figure 1B; Dataset S2). Reduction of total photosynthetic activity is a common feature of plant stress, and the results confirm that the ability to maintain the photosynthetic capacity is a good indicator of tolerance to either *neuSAL* or *alkSAL* in *A. thaliana.* These observations point to a better performance of demes from pre litoral calcareous soils (V1^PC^, LM2^PC^) under *alkSAL* in terms of growth maintenance and photosynthesis, and agree with the severe reduction in rosette diameter previously reported on coastal demes under *alkSAL* (Pérez-Martín et al., 2022).

**Figure 1.**
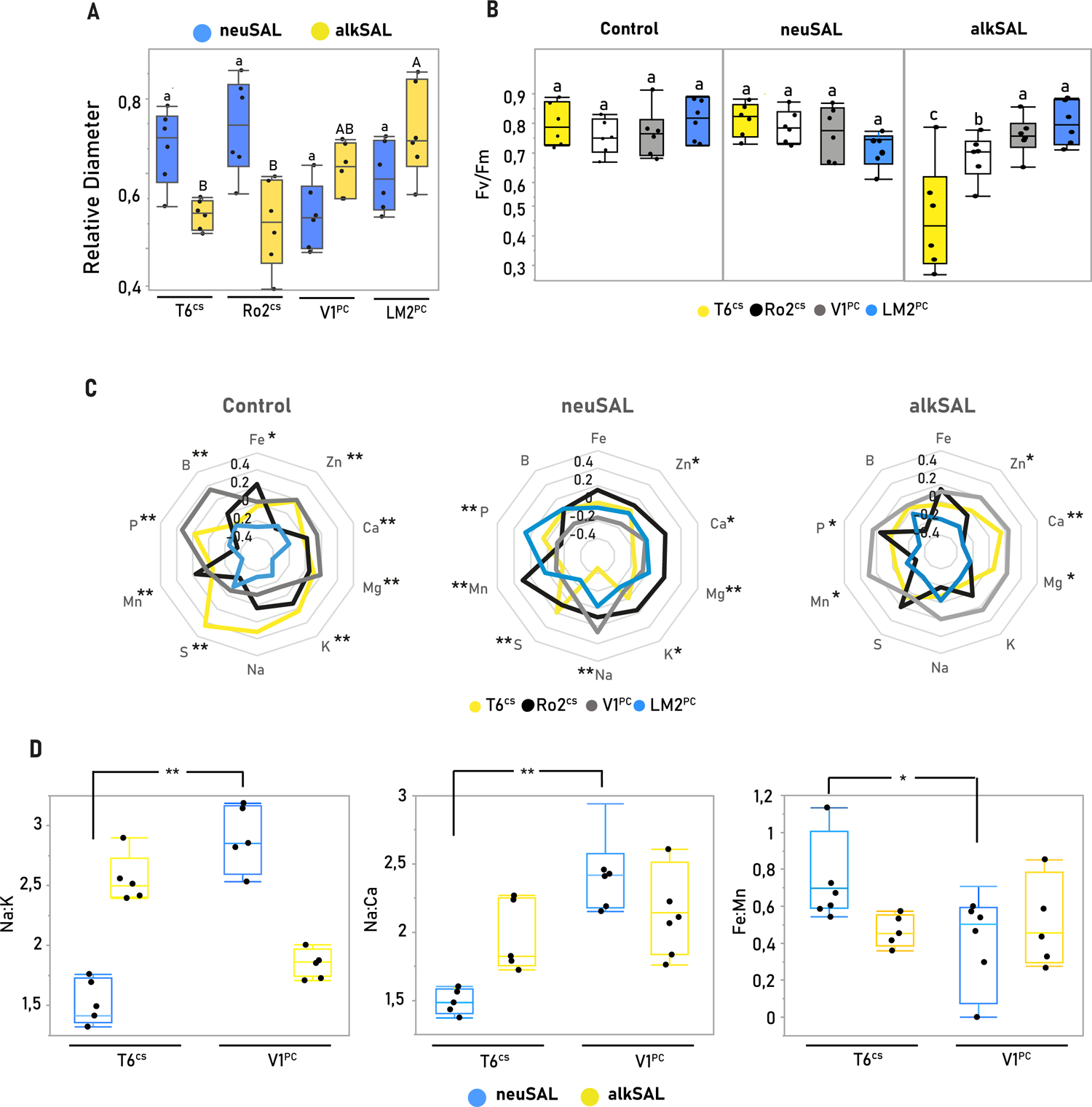
Leaf growth, photosynthesis efficiency and mineral nutrition in *A. thaliana* demes under control (C), neutral (*neuSAL*) and alkaline (*alkSAL*) salinity. (A) Mean ± SE of growth (relative rosette diameter) of each deme and treatment. Each dot represents a biological replicate (n=6). Mean values with different letters indicate significant differences (Tukey’s HSD, adj. *p*-value < 0.05). **(B)** Mean ± SE of Maximum PSII photochemical efficiency (Fv/Fm) of each deme and treatment. Each dot represents the mean value of 2 technical replicates (2 measurements on each biological replicate) (n=6). Mean values with different letters indicate significant differences (Tukey’s HSD, adj. *p*-value < 0.05). **(C)** Radial plots showing normalized differences of 10 elements in plant leaf samples from each deme and treatment. Axes display *Z-*scores calculated per element, and elements exhibiting significant differences in at least 1 deme (Tukey’s HSD, adj. *p*-value < 0.05) are marked with an asterisk on each radial plot. **(D)** Mean ± SE of analyzed nutrient ratios in the two contrasted demes T6^CS^ and V1^PC^. Each dot represents a biological replicate (n=6). Nutrient ratios exhibiting significant differences for each deme between salinity treatments (Student *t*-Test, adj. *p*-value < 0.05) are marked with an asterisk (*: adj. *p*-value < 0.05; **: adj. *p*-value < 0.01). Plants were grown in potting mix soil and irrigated with C (0.5-Hoagland, pH 5.9), *neuSAL* (100 mM NaCl, pH 5.9) or *alkSAL* (85 mM NaCl, 15 mM NaHCO3, pH 8.3) for 2 weeks.

#### K^+^ and Fe uptake efficiency are indicators of plant tolerance to salinity

Mean-standardized values (– 1< value >1) of leaf and soil elemental contents were used for the analysis. Overall, significantly differentiated leaf nutrient profiles (*p*-value ≤ 0.05, Tukey’s HSD) were observed for all demes in most elements under control, *neuSAL* and *alkSAL* treatments (Figure 1C; Datasets S3). Whereas nutrient profiles were clearly genotype-specific under control conditions, *neuSAL* highlighted a significantly higher K^+^ levels in T6^CS^ and Ro2^CS^ when compared to V1^PC^ and LM2^PC^, on one hand, and opposite trends in leaf Na^+^ accumulation in coastal demes, on the other: T6^CS^ accumulated significantly less leaf Na^+^ when compared to all other demes, but Ro2^CS^ increased leaf Na^+^ due to reported allelic variation at the HKT1;1 locus, which leads to higher shoot Na^+^ accumulation in individuals bearing the weak *HKT1;1^HLS^* allele (Rus et al., 2006; Busoms et al., 2018). Still, highest leaf Na^+^ accumulation under neuSAL was reported in V1^PC^ (*p-*value = 1,40E-04; Tukey’s HSD). Under *alkSAL*, V1^PC^ outrated all other demes in leaf nutritional status, while LM2^PC^ displayed a general significant reduction in leaf nutrient concentration.

V1^PC^ and LM2^PC^ have been previously classified as good performers under *alkSAL* based on growth, reproductive and physiological traits when compared to coastal T6^CS^ and Ro2^CS^, and the most tolerance phenotypes under neutral and alkaline salinity were observed for T6 and V1, respectively (Pérez-Martín et al., 2022). Therefore, nutrient ratios with reported biological importance under salinity conditions were compared in T6^CS^ – as tolerant to *neuSAL* but sensitive to *alkSAL* – and V1^PC^ – as sensitive to *neuSAL* but good performer under *alkSAL*, to further explain the implications of the differential nutrient accumulation under each type of salinity (Figure 1D; Dataset S4). Na:K ratio in T6^CS^ (1.51) was significantly lower (*p*-value ≤ 0.05, Tukey’s HSD) than V1^PC^ (2.88) under *neuSAL*, but higher (2.56 and 1.86 for T6^CS^ and V1^PC^, respectively) under *alkSAL*. It has been previously concluded that salt tolerance is driven more by a plants ability to retain K^+^ under salt stress, than its ability to simply exclude Na^+^ ions (Wang et al., 2007; Shabala & Cuin, 2008). The Na:Ca ratio followed the same trend as Na:K, being significantly lower in T6^CS^ (1.49) than in V1^PC^ (2.88) under *neuSAL*. Cellular Na^+^ and Ca cell transport pathways are linked via SOS2 (a ser/thr protein kinase activated by SOS3, a calcium sensor protein), as SOS2 regulates SOS1 and NHX1 (plasma membrane and tonoplast Na^+^/H^+^ antiporters) but also CAX1 (a vacuolar Ca/H^+^ antiporter) (Cheng et al., 2004). Decreasing Na:Ca ratio is shown to enhance K^+^ uptake and reduce Na^+^ uptake, ultimately decreasing Na:K (Garg et al., 2015). This seems to be the case for V1^PC^ under *alkSAL* but not under *neuSAL*, which might be causing its sensitivity to neutral but not to alkaline salinity. Moreover, Fe uptake efficiency is critical in a context of alkalinity, as high pH decreases Fe availability and leads to plant chlorosis (Hsieh et al., 2016). Under *neuSAL*, the Fe:Mn ratio was increased in T6^CS^ (0.77) due to a higher leaf Fe accumulation when compared to V1^PC^ (0.01 vs −0.20). Under *alkSAL*, the maintenance of Fe:Mn despite the high increase in leaf Mn concentration displayed by V1^PC^ (0.30) indicates higher Fe uptake or translocation efficiency (Figure 1D). This suggests that T6^CS^ – adapted to moderate levels of neutral salinity - could be hampered by the alkalinity component of *alkSAL*. Overall, the observed response of each deme to *alkSAL* seems driven by other factors than the Na^+^ content in their native soils, as it is the case for *neuSAL* (Busoms et al., 2018), and provides evidence that tolerance to salinity in siliceous soils does not confer tolerance to alkaline salinity (Pérez-Martín et al., 2022).

#### Differential elicitation of phytohormonal signaling pathways depends on salinity type and deme

Considering that regulation of phytohormone levels is one of the most important mechanisms conferring tolerance to a wide spectrum of abiotic stresses (Zhang et al., 2019), endogenous phytohormones were quantified in leaf tissue of the studied demes. Average concentrations of the analyzed phytohormones can be found in Dataset S5. A significant elicitation (*p*-value ≤ 0.01, Student’s *t-*Test) of IAA, ACC, and JA was observed under *alkSAL* when compared to control and *neuSAL* treatment (Figure 2 A-C), and a decrease in leaf SA and an increase in leaf ABA concentrations were caused by both *neuSAL* and *alkSAL* conditions (Figure 2 D-E), which supports previous observations on ABA-SA and JA-SA antagonistic signaling (Kazan & Manners, 2012; Moeder et al., 2010). In addition, the studied demes exhibited an origin-dependent accumulation profile for ABA under *neuSAL*, by which coastal individuals (T6^CS^, Ro2^CS^) displayed higher ABA accumulation when compared to non-coastal V1^PC^ and LM2^PC^ (Figure 2 E). Salinity-induced ABA biosynthesis causing stomatal closure, reducing stomatal conductance and water consumption, and promoting overall plant tolerance to salinity is well reported (Lovelli et al., 2012). Here, alkaline salinity elicits higher alteration of hormonal networks than neutral salinity, whereas ABA accumulation seems to contribute to the enhanced response of demes locally adapted to *neuSAL* conditions by increasing water use efficiency (Cardoso et al., 2020) and restricting K^+^ efflux (Chen et al., 2022). In fact, ABA concentration in coastal demes under *neuSAL* correlated with increased plant relative water content (RWC) (Figure 2 F), suggesting that ABA enhances leaf salt resistance by improving the plant water status through reduced stomatal conductance.

**Figure 2.**
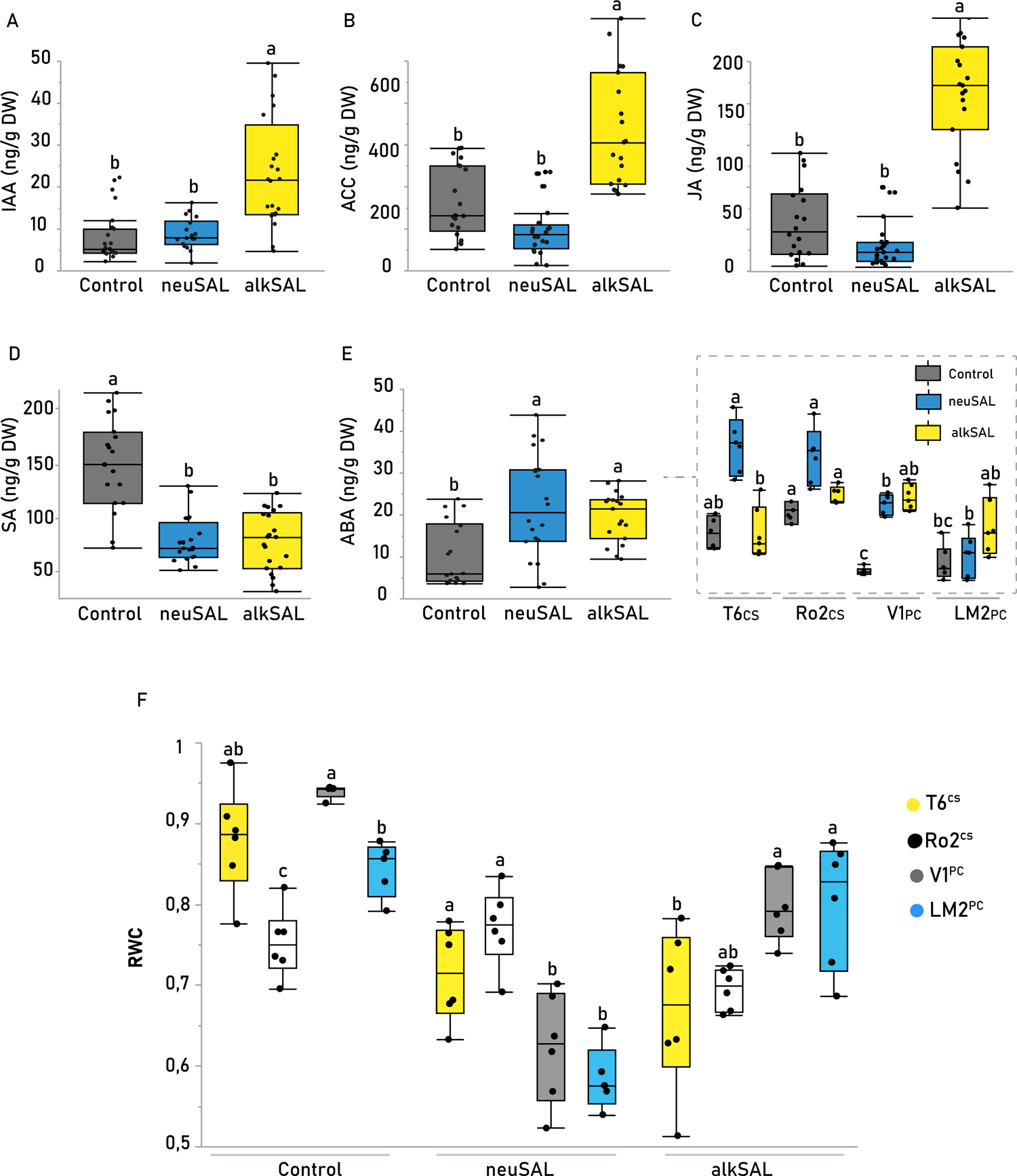
Endogenous phytohormone content in *A.thaliana* demes under control (C), neutral (*neuSAL*) and alkaline (*alkSAL*) salinity. Figure 2. Endogenous phytohormone content in A.thaliana demes under neutral (neuSAL) and alkaline (alkSAL) salinity. Mean ± SE of relative (A) Indole acetic acid (IAA), (B) 1-Aminocyclopropane-1-carboxylic acid (ACC), (C) Jasmonic Acid (JA), (D) Salicylic Acid (SA) and (E) Abscisic Acid (ABA) concentration in all demes under control and salinity treatments. Each dot represents a biological replicate (n=6), and treatments are color-coded (see legend). Zoomed boxplots show relative phytohormone concentration for each deme. Mean values with different letters indicate significant differences (Tukey’s HSD, adj. p-value < 0.05). Plants were grown in potting mix soil and irrigated with C (0.5-Hoagland, pH 5.9), neuSAL (100 mM NaCl, pH 5.9) or alkSAL (85 mM NaCl, 15 mM NaHCO3, pH 8.3) for 2 weeks.

#### Multivariate analyses reveal new players defining the alkalinity component of salinity

Multivariate analyses on all measured traits were performed for all samples by means of Principal Component (PCA) and Multiple Pearson Correlation (MPC) analyses. The PCA showed that eigenvalues of the two first principal components represented up to 48% (PC1 25%; PC2 23%) of the total variance (*p-*value = 1.13E-09 for PC1 and 0.0002 for PC2 (Dataset S6). A general look at the PC1–PC2 axes plane plot (Fig 3A) shows that alkalinity contribute to the construction of component 1 (samples submitted to neutral or alkaline salinity differentiate along the x-axis), whereas salinity contributes to PC2 construction (control samples differentiate from *neuSAL* and *alkSAL* samples along the y-axis). The loading matrix revealed that well correlated mineral nutrient traits (leaf Zn and leaf Mg), as well as endogenous JA, were the variables contributing to PC1 with a loading factor above 0,70 (Dataset S7). In fact, *alkSAL* samples contained significantly higher Zn than *neuSAL* samples (*p*-value ≤ 0.05, Tukey’s HSD) (Figure 3B). In turn, MPC results showed that Zn displayed the highest correlation with Mg and JA under *alkSAL* (R^2^ = 0.8; R^2^ = 0.6), but correlation with Mg was lost (R^2^ = 0.15), and correlation with JA was inverted (R^2^ = −0.48) under *neuSAL* (Figure 3C).

**Figure 3.**
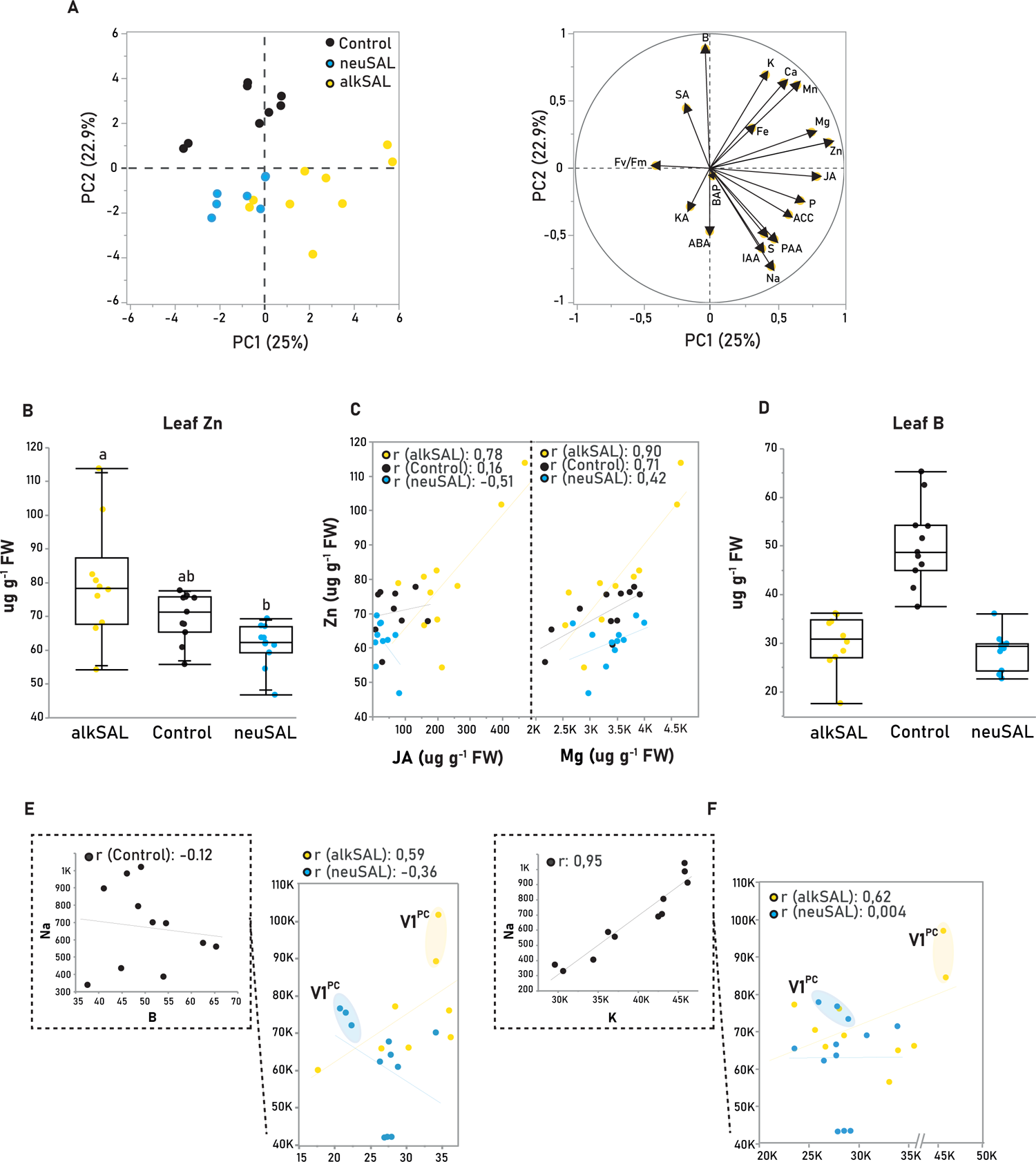
Multivariate analysis on physiological traits in *A. thaliana* demes under control (C), neutral (*neuSAL*) and alkaline (*alkSAL*) salinity. (A): Principal Component Analysis (PCA). Left: individual factor map with treatment-colored samples distribution on the 2 most informative axes (PC1 and PC2) of the PCA. Right: variable factor map presenting the amount of variance from each trait on the total variance in the PCA. Longer arrows represent larger amount from total variance. **(B)** Mean ± SE of leaf Zn (µg g^-1^ DW) in all demes under control and salinity treatments. **(C)** Scatter plot matrix section from multivariate correlation analysis of all physiological traits. Scatter plots show correlation between leaf Zn [(µg g^-1^ DW), y-axis] and leaf JA and Mg [(ng g^-1^ FW), left; (µg g^-1^ DW), right; x-axis] in all samples. **(D)** Mean ± SE of leaf B (µg g^-1^ DW) in all demes under control and salinity treatments. **(E)** Scatter plot matrix section from multivariate correlation analysis of all physiological traits. Scatter plots show correlation between leaf Na [(µg g^-1^ DW), y-axis] and leaf B (left) and K (right) (µg g^-1^ DW); x-axis] in all samples. r = correlation coefficient. Each dot represents a biological replicate (n=3 per deme) and treatments are color-coded (see legend). Letters indicate significant differences (Tukey’s HSD, adj. *p*-value < 0.05). Plants were grown in potting mix soil and irrigated with C (0.5-Hoagland, pH 5.9), *neuSAL* (100 mM NaCl, pH 5.9) or *alkSAL* (85 mM NaCl, 15 mM NaHCO3, pH 8.3) for 2 weeks.

Leaf Na^+^, leaf K^+^ and leaf B contributed to the construction of component 2, being leaf B dramatically decreased in the presence of salinity regardless of neutral or alkaline (Figure 3D). MPC showed how Na-B correlated negatively under *neuSAL* (R^2^ = −0.42) but positively under alkSAL (R^2^ = 0.35), whereas no correlation was found in control (R^2^ = 0.03) (Figure 3E). While co-toxicity of salinity and B is reported (Pandey et al., 2019), few studies on neutral or alkaline salinity causing B deficiency are found. Boron alleviates salinity and saline-sodicity stress by improving leaf water and soluble carbohydrate contents as well as maintaining osmotic potential (Yousefi et al., 2020; Mehmood et al., 2009). Remarkably, V1 displayed the highest B and K^+^ maintenance under *alkSAL* and the highest B and K^+^ deficiency under *neuSAL* (Figure 3E-F).

In summary, the reported results suggest that growth and PSII efficiency maintenance are indicators of differential tolerance to neutral and to alkaline salinity in the studied demes. Whereas neutral salinity tolerance in coastal individuals is driven by a decreased Na^+^ shoot translocation in T6^CS^ and by the ability to retain K^+^ in Ro2^CS^, the maintenance of leaf K^+^ homeostasis by increasing leaf Ca^2+^ accumulation is a requirement for a better response to alkaline salinity. Moreover, enhanced tolerance to *alkSAL* in V1^PC^ is partially explained by higher Fe, B and K^+^ internal use efficiency. Differential modulation of auxin, ABA and CKs among genotypes is likely involved in tolerance to either *neuSAL* or *alkSAL* in the studied demes; and differences in leaf Zn concentration under *neuSAL* and *alkSAL* may affect endogenous phytohormone content through the stimulation of tryptophan biosynthesis, which is a precursor of indoleacetic acid and a potential conjugate of IAA and JA.

### 2. Transcriptome profiling of A. thaliana under neutral and alkaline salinity

Leaf transcriptome profiling of the 4 demes cultivated under control, *neuSAL* or *alkSAL* conditions was performed to characterize the mechanisms underlying the differential response to neutral and alkaline salinity at the gene expression level. Multi-Dimensional Scaling (MDS) was used to visualize global transcriptional similarity among samples (Figure 4A). The transcriptome of each sample was shaped by deme and colored by treatment. Accordingly, individuals showed higher transcriptional similarity between samples corresponding to the same treatment, indicating that the studied transcriptomic response is treatment specific.

**Figure 4.**
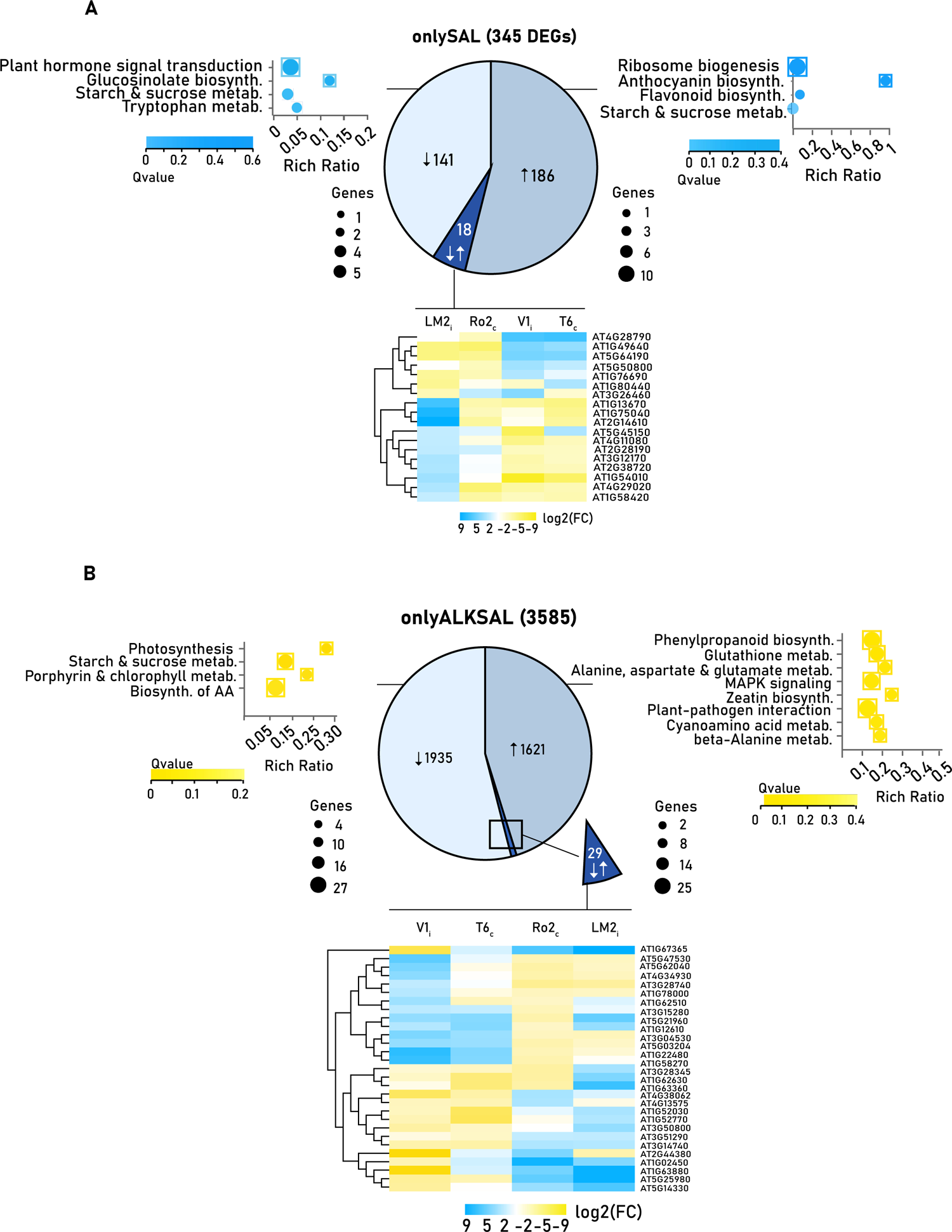
Differential Gene Expression (DEG) analysis in *A. thaliana* demes T6^CS^ (yellow), Ro2^CS^ (black), V1^PC^ (blue), and LM2^PC^ (grey). (A) Multidimensional scaling (MDS) plot of RNA expression profiles in the studied samples. Sample relations are plotted using a multidimensional scaling plot (MDS) generated with edgeR showing the variability between replicates and treatments in log2-fold-change distance. The axes represent gene expression levels between the different experimental factors. Each dot represented a sample, color coded by treatment (Black: Control, Blue: *neuSAL*, Yellow: *AlkSAL*) and shape coded by deme (Color-filled square: T6^CS^, empty square: Ro2^CS^, color-filled octagon: V1^PC^, empty octagon: LM2^PC^). **(B)** Histogram showing the number of differentially expressed genes for each deme (n=3) and treatment (*neuSAL*/Control; *alkSAL*/Control). **(C)** Venn diagram showing the mutual overlaps of DEGs among demes under each treatment. DEGs were filtered at log fold change (LFC > 2, LFC < −2), and *q*-value (FDR-adjusted *p*-value) < 0.05.

Differentially expressed genes (DEGs) per deme and treatment were declared based on absolute log_2_ fold-change (FC) > 2 under each treatment compared to control conditions (*neuSAL*/Control, *alkSAL*/Control) (*q-*value < 0.05, Wald test). When considering all demes, a total of 6164 DEGs (2559 up-regulated and 3605 down-regulated) were identified under neuSal and 10487 DEGs (4995 up-regulated and 5492 down-regulated) under *alkSAL*. A 65% of total DEGs from *neuSAL* were found in T6^CS^ and V1^PC^, while 29% and 34% in Ro2^CS^ and LM2^PC^. Under *alkSAL*, T6^CS^, Ro2^CS^, V1^PC^ and LM2^PC^ displayed 1.9, 4.1, 2.2 and 1.53-times more DEGs than under *neuSAL*. Under *alkSAL*, T6^CS^, still displayed the most altered transcriptome, followed by Ro2^CS^ (73% and 68% of total *alkSAL* DEGs, respectively), while V1^PC^ and LM2^PC^ transcriptomic response was more attenuated (58% and 44% of total DEGs) (Figure 4B-C).

#### Exclusive and shared molecular components of neutral and alkaline salinity

To gain more insight into commonalities and particularities between treatments, the subset of DEGs shared by at least 3 out of 4 demes in *neuSAL* and *alkSAL* were selected and merged, thus obtaining individual and common DEGs under both treatments (Figure 4D). On the one hand, 345 of DEGs were found in response to *neuSAL* but not to *alkSAL* and were referred as *onlyNEUSAL* DEGs set (Table S1). From these, 141 and 186 (41% and 54% of transcripts, respectively) were commonly upregulated or downregulated among demes, and 18 (5%) showed ambiguous expression trends depending on the deme (Figure 5A). On the other hand, 3585 DEGs changed under *alkSAL* but not under *neuSAL* and were referred to as *onlyALKSAL* DEGs set (Table S2). From these, 1621 and 1935 (45% and 54% of transcripts, respectively) were upregulated or downregulated, and only a minimal fraction of 29 (0,8%) showed opposite expression trends in at least 1 deme (Figure 5B). Finally, the 1219 of the selected DEGs were shared under both *neuSAL* and *alkSAL* and were referred as *coreSAL* DEGs set (Table S3). From these, 406 (65%) were upregulated, 786 (33%) downregulated and 27 (2%) with opposite expression trends in at least one deme (Figure 7A). It is well known that plants facing complex stress conditions trigger cascade of signals unique to specific stress components as well as shared responses (Zandalinas et al., 2020; Skalak et al., 2021). Studies assessing *Arabidopsis* response to combinatorial stresses described antagonistic expression trends for only 5% to 10% of transcripts, which is in line with the expression trends found for each DEGs subset described above. Moreover, 86% of DEGs from the *coreSAL* were found in the 4 studied demes (Figure S3), which reveals that contribution to the *coreSAL* DEGs set is high for all studied demes and, thus, similar strategies in the studied individuals lead to differential responses to *neuSAL* and *alkSAL*.

**Figure 5.**
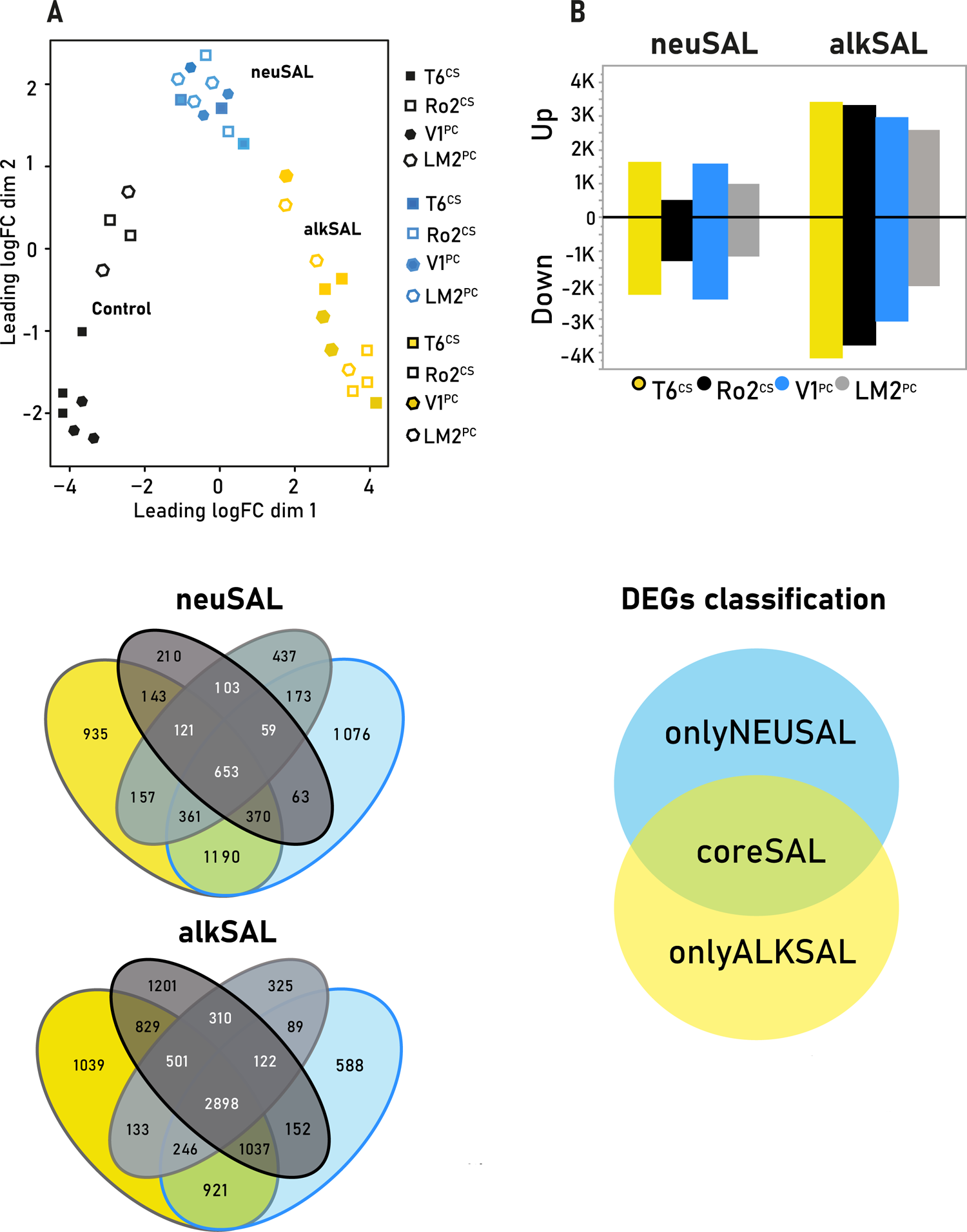
Characterization of the onlySAL and onlyALKSAL sets. DEGs were filtered at log fold change (LFC > 2, LFC < −2), and q-value (FDR-adjusted p-value) < 0.05. Chart pie showing expression trend of total **(A)** onlySAL, **(B)** onlyALKSAL and **(C)** coreSAL DEGs. KEGG pathway enrichment analyses of downregulated (bubble plot, left) and upregulated DEGs (bubble plot, right) are shown for each corresponding DEG set. In bubble plots, pathways belonging to different classifications are listed on the left. Only the top 4 are presented, which are sorted by the q-value. Rich ratio (x-axis) is the ratio of the DEG number to the total gene number in a certain pathway. Bubble size represents number of DEGs included in each pathway (see legend) and enclosed bubbles indicate significant pathway enrichment. In heatmap: genes were clustered according to their expression pattern (Euclidean distance) and column clustering groups samples based on gene expression similarity. Color scale indicates the expression levels (yellow, low expression; blue, high expression).

The overall results obtained from DEG analysis and classification provide evidence that the stress imposed on plants by alkaline salinity is distinct from that imposed by salinity in a neutral soil (Guo et al., 2010) and that alkaline salinity involves more complex processes than neutral salinity et the transcriptomic level (Fang et al., 2021).

#### KEGG Erichment analysis of onlyNEUSAL and onlyALKSAL DEGs

Kyoto Encyclopedia of Genes and Genomes (KEGG) pathway analysis was performed in *onlyNEUSAL* and *onlyALKSAL* DEGs to identify which categories were significantly enriched in metabolic pathways (*q*-value < 0.05), and thus target the presence of alkalinity-dependent salinity effects in the studied demes. *onlyNEUSAL* and *onlyALKSAL* enriched pathways are listed in Table S4.

#### Upregulated transcripts in onlyNEUSAL and onlyALKSAL reveal higher demand for antioxidant protection under alkaline salinity

In *onlyNEUSAL*, only 2 pathways containing 6 and 2 upregulated transcripts were significantly enriched: “ribosome biogenesis” and “anthocyanin biosynthesis”. This could indicate that expression of such transcripts is triggered by the NaCl component, likely Cl^-^ ion toxicity, which is present in higher extent under *neuSAL* than under *alkSAL* is reported to be the most significant toxic component of the saline solution in some species (Munns & Tester, 2008) and, more specifically, strongly decreases ribosome stability (Brady et al., 1984). The role of anthocyanin accumulation in the positive regulation of tolerance mechanisms to abiotic factors is well reviewed (Perea-Resa et al., 2017; Naing & Kim, 2021; Kubra et al., 2021; Zhang et al., 2019), among others, has been reported. Here, activation of anthocyanin biosynthesis genes (AT4G14090 and AT5G54060) could take place for scavenging ROS excess due to plant oxidative damage (Naing & Kim, 2021). In *onlyALKSAL*, the most enriched pathways with up-regulated transcripts were phenylpropanoid biosynthesis (25 DEGs) and glutathione metabolism (18 DEGs) (Figure S4). Phenylpropanoid metabolism is elicited by abiotic stresses such as drought and salinity and, moreover, is reported to contribute to salinity-alkalinity tolerance (Gan et al., 2021). The pH-dependent increase in expression of DEGs from phenylpropanoid and glutathione signaling pathways is in line with that reported in recent studies (Waters et al., 2018, Gan et al., 2021; Han et al., 2018).

The above results point to a pH-dependent response in the studied demes and a higher requirement of the plant antioxidant machinery in the presence of alkaline salinity, which would lead to an enhanced activation of transcripts from the phenylpropanoid and glutathione metabolism.

#### Salinity-triggered phytohormone responses are alkalinity-dependent

Endogenous phytohormone quantification of the studied demes under *neuSAL* and *alkSAL* treatments revealed a sustained biosynthesis of auxins, ACC and JA under prolonged plant exposure to alkaline salinity. In turn, higher ABA after 2 weeks of *neuSAL* irrigation was observed only on genotypes locally adapted to neutral salinity (Figure 2). To identify activation or repression patterns in signaling pathways accounting for the differential phytohormone profiles, the expression levels of phytohormone-responsive genes under each treatment were analyzed (*q-*value < 0.05; Student *t*-Test) (Dataset 13). A general trend towards gene repression was present (Figure 6), as previously reported for other abiotic stress long term transcriptional responses (Leyva-Pérez et al., 2015).

**Figure 6.**
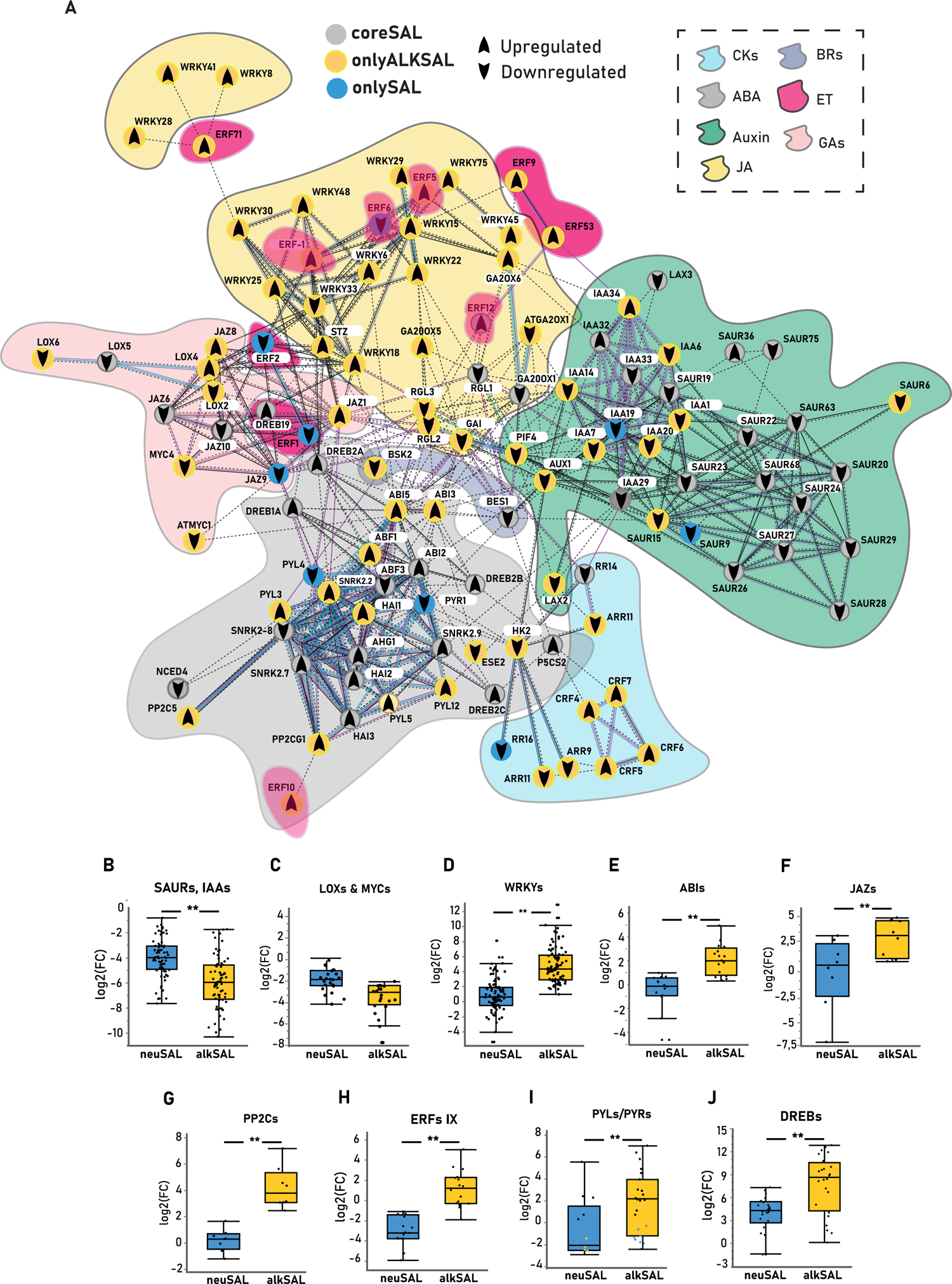
Global overview of phytohormone signaling dynamics. (A) Protein-protein interaction (PPI) network of the analyzed DEGs involved in phytohormone signaling based on the STRING online database (https://string-db.org/). Nodes are color-coded according to each DEGs subset, arrows inside nodules indicate expression trend and colored surrounding areas distinguish between different hormone classes (see legends). (B) Expression levels [mean ± SE log2(FC)] of DEGs from (B) SAURs & IAAs (n=56), (C) LOXs/MYCs (n=20), (D) WRKYs (n=44), (E) ABIs (n=16), (F) JAZs (n=8), (G) PP2Cs (n=8), (H) IX clade ERFs (n=12), (I) PYLs/PYRs (n=23) and (J) DREBs (n=24) in all demes among treatments. Asterisks indicate significant differences in mean expression values among treatments (Student t-test, adj. p-value < 0.05; *: adj. p-value < 0.05; **: adj. p-value < 0.01). Neutral salinity: *neuSAL*; alkaline salinity: *alkSAL*.

**Figure 7.**
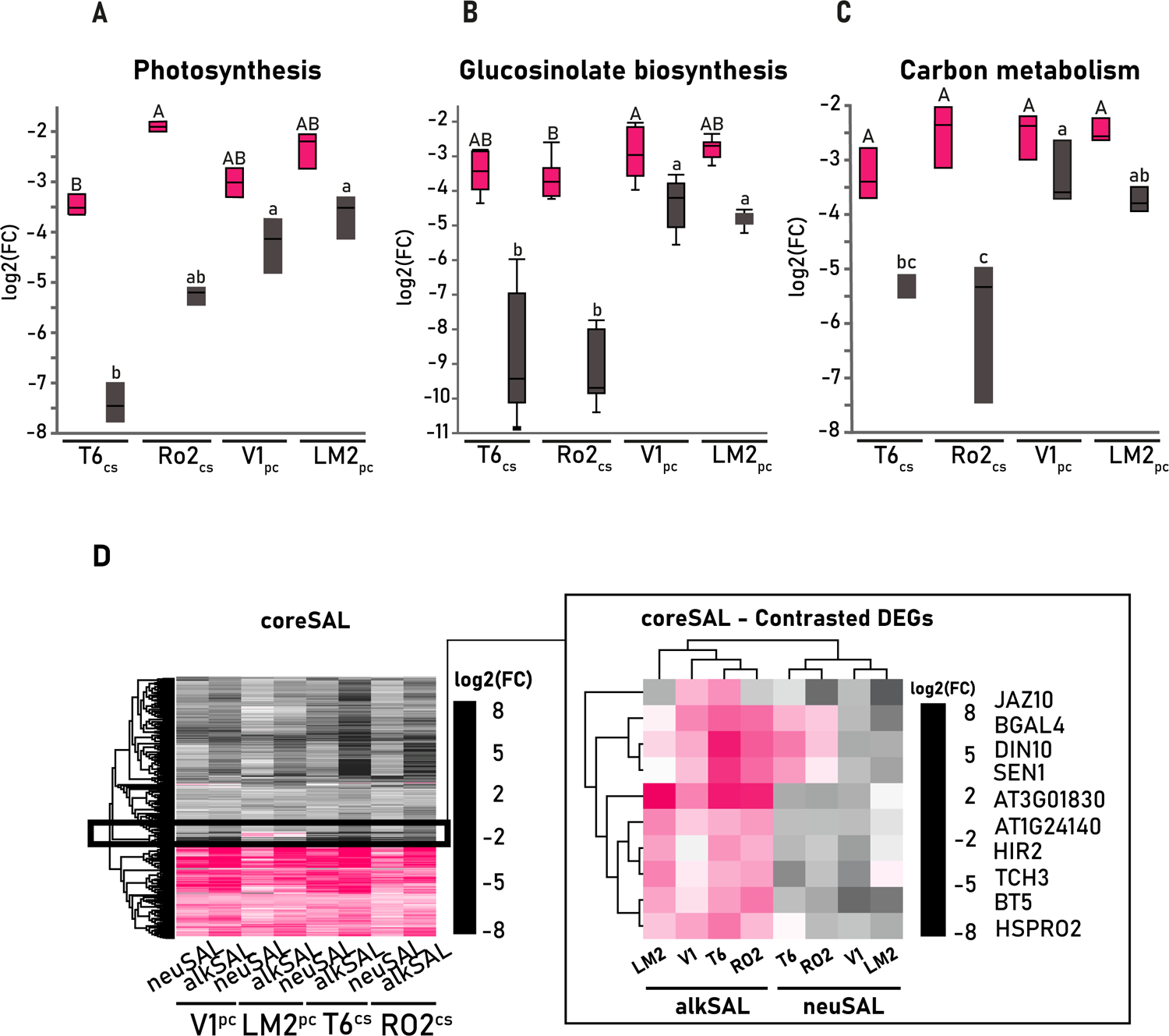
Characterization of the coreSAL DEGs set. DEGs were filtered at log fold change (LFC > 2, LFC < −2), and *q*-value (FDR-adjusted *p*-value) < 0.05. Expression levels [mean ± SE log2(FC)] of **(A)** “photosynthesis”, **(B)** “glucosinolate metabolism” and **(C)** “carbon metabolism” DEGs in each deme (n=3) and treatment. Letters indicate significant differences in mean expression values among demes (Tukey’s HSD, adj. *p*-value < 0.05). **(D)** Heat map showing the expression profile of coreSAL DEGs (1219 genes) for each deme between neutral and alkaline salinity when compared to control (neuSAL/C; alkSAL/C). Genes were clustered according to their expression pattern (Euclidean distance). In frame, heatmap showing the expression profiles of coreSAL DEGs with contrasted expression trends among salinity types and/or demes. The color scale indicates the expression levels (grey, low expression; pink, high expression). Genes were clustered according to their expression pattern (Euclidean distance), while column clustering groups samples based on gene expression similarity.

Higher inhibition of auxin (SAURs, Aux/IAAs) and JA (LOX, MYCs) responsive transcripts was observed after 2 weeks of *alkSAL* exposure when compared to neutral salinity (Figure 6 B-C). In contrast, *alkSAL* treatment caused an induction of the transcription factor WRKY and ABI (*ABA-insensitive*) families, of repressors of the JA pathway (JAZs), and of protein phosphatases activating abscisic acid-inducible gene expression (PP2C5 & PP2CG1), whereas their average expression levels showed to be attenuated after prolonged exposure to *neuSAL* (Figure 6 D-G). Opposite expression trends were displayed among treatments in ethylene and ABA responsive transcripts (ERFs and PYL/PYRs), in which significant activation under *alkSAL* but inhibition under *neuSAL* were observed (Figure 6 H-I). Finally, DREBs (*Dehydration responsive element binding*) showed sustained activation under both *neuSAL* and *alkSAL,* with significantly higher transcript expression under *alkSAL* (Figure J).

The downregulation of SAURs during the late phase of the plant salinity response agrees with a protein degradation targeted after the observed auxin accumulation (Figure 2). Meanwhile, the reduced expression of the early auxin-responsive SAURs and the auxin co-receptors Aux/IAAs under *alkSAL* suggest further repression of auxin signaling mediated by the presence of JA and ABA rather than the removal of the inducer (IAA) (Markakis et al., 2013; van).

Under alkaline salinity, an antagonistic interaction between ABA and JA signaling pathways modulating stress-responsive gene expression (Anderson et al., 2004; Kazan & Manners, 2013; Sah et al., 2016) is revealed, as depicted by the significant upregulation of ABA receptors (PYLs), ABA responsive genes (ABIs) and other ABA-dependent MAPK phosphatases (PP2Cs), and the enhanced inhibition of JAs responsive transcripts (LOXs and MYCs) in *onlyALKSAL*. Moreover, the expression of JAZs, targeting MYCs repression, was only induced by *alkSAL.* This supports a negative regulatory feedback loop occurring after long exposure to alkaline salinity to turn off the transcriptional reprogramming caused by JA accumulation under *alkSAL* and to minimize JA inhibition on plant growth, in an attempt of acclimation (Riemann et al., 2015). In turn, the stimulation of ABA signaling and accumulation, and the NaCl tolerance observed in coastal demes under *neuSAL* are likely mediated by WRKYs (Hussain et al., 2021), which remained active after 2 weeks of *neuSAL* treatment. Among them, enhancers of the osmotic, ROS and hormone signaling pathways under salinity stress have been recently reported (Price et al., 2022).

Most DREBs regulate the expression of multiple stress-inducible genes and play pivotal roles in the plant abiotic stress response. All identified DREBs were strongly upregulated after 2-week exposure to both *neuSAL* and *alkSAL.* Among them, DREB1A and DREB2C, reported as two of the DREBs containing the most types of abiotic stress-related motifs in the promoter region (Xie et al., 2019), were found. Upregulated DREBs in the plant long-term *coreSAL* gene set provide further evidence that DREBs form a central network to control the plant salinity response also at the late stage (Sazegari et al., 2015). Contrarily, sustained upregulation of twelve DEGs from another subfamily of AP2/ERF transcription factors, ERF-IX, seems to be required solely by alkaline salinity stress and associates with the plant significant ACC accumulation under *alkSAL* (Wang etal., 2022).

Overall, differences in the transcriptional regulation of phytohormones-related genes under prolonged plant exposure to either neutral or alkaline salinity were observed. The above-mentioned expression patterns are consistent with a higher alteration in the plant transcriptome caused by alkaline salinity (Figure 4B) leading to a more extensive rearrangement in the phytohormonal signaling network. In this regard, transcript inhibition for protein turnover seems to be a regulation mechanism for phytohormone signaling pathways such as auxin and JA under *alkSAL,* and ABA under *neuSAL* (McClellan & Chang, 2008). An implication of differential auxin and ABA regulation mechanisms for proton extrusion, through H^+^-ATPase activation by inhibition of PP2C proteins should be further addressed, as it is an essential Fe-deficiency plant response (Sun et al., 2020; Spartz et al., 2014).

#### KEGG Erichment analysis of coreSAL DEGs

KEGG pathway analysis was performed in *coreSAL* DEGs to identify significantly enriched metabolic pathways (*q*-value < 0.05) in the general salinity stress response and the presence of genotype-dependent salinity effects in the studied demes. *coreSAL* enriched pathways are listed in Table S6.

#### coreSAL transcript activation reflects shared salinity stress responses

The significantly enriched pathways in the *coreSAL* set were “Ribosome biogenesis in eukaryotes”, “Protein processing in endoplasmic reticulum” and “Cutin, suberin and wax biosynthesis” (Table S6). Ribosome biogenesis and protein synthesis in the ER are the two pathways comprising cellular processes with highest metabolic cost (Piazzi et al., 2019). Here, we report a consistent upregulation of genes involved in pre rRNA and RNA processing and maturation (FIB2, PWP2, AT4G04940, AT1G67120, AT3G06530), nucleolytic cleavage (AT5G22100), and cell differentiation (AT2G18900) in all 4 demes under neutral and alkaline salinity, followed by the activation of 11 Heat Shock Proteins – which encode for molecular chaperones that interact with ribosomes ensuring their stability (Piazzi et al., 2019) (Figure S5A). This indicates nucleolar stress (Yang et al., 2018) triggered by both neuSAL and alkSAL that leads to the activation of the protein synthetic machinery in the studied demes (Figure S5B). Moreover, enhanced activation of 2 genes involved in suberin deposition (FAR1, FAR5) – previously reported to be transcriptionally induced by salinity (Domergue et al., 2010) - were observed in all demes under *neuSAL* and *alkSAL* (Figure SC), which suggests that increase of suberin deposition may enhance tolerance to neutral and alkaline salinity (Zhang et al., 2021).

#### coreSAL transcript inhibition reveals differential sensitivities to alkSAL in the studied demes

The significantly enriched pathways containing down-regulated transcripts detected in the *coreSAL* set were ‘Photosynthesis’ and ‘Photosynthesis antenna proteins’, ‘Glucosinolate biosynthesis’ ‘Glyoxylate and dicarboxylate metabolism’, ‘Carbon fixation’, ‘Carbon metabolism’, ‘2-Oxocarboxylic acid metabolism’, ‘Nitrogen metabolism’, and ‘Glycine, serine and threonine metabolism’ (Figure 5C; Table S6). T6^CS^ suffered the highest inhibition of all DEGs from “Photosynthesis” and “photosynthesis antenna proteins” (an average of 1.5 and x1.8 times more downregulation than V1^PC^ and LM2^PC^, respectively), followed by Ro2^CS^ in 2 transcripts: PSAH1-2 (an average of x1.5 times more downregulation in T6^CS^, Ro2^CS^ when compared to V1^PC^ and LM2^PC^ under *alkSAL*) (Figure 7A). PSAH1 and PSAH2 encode subunits H of photosystem I reaction center and their up-regulation under neutral salinity has been reported in halophytes (Hao et al., 2020). The downregulation of photosynthesis-related transcripts in the *coreSAL* response agrees with previous transcriptome studies under neutral salinity reporting a correlation between down regulation of antenna proteins and higher salt sensitivity (Guo et al., 2017; Jing et al., 2019). Similarly, DEGs from “glucosinolate biosynthesis” and a DEG subset from “carbon metabolism” displayed an average of x1.7 times more downregulation in T6^CS^ and Ro2^CS^ when compared to V1^PC^ and LM2^PC^ under *alkSAL* (Figure 7B).

Glucosinolates are secondary metabolites mainly found in Brassicaceae. Downregulation of genes involved in glucosinolates biosynthesis has been reported under several types of salts (Aghajanzadeh et al., 2017). A previous study (Pérez-Martín et al., 2021) characterized the response to bicarbonate exposure of T6^CS^ and A1_i_ – as a carbonate sensitive and carbonate tolerant deme, respectively – and reported the downregulation of several GSTU genes involved in glucosinolate metabolism only in T6^CS^. Two Ribulose Biphosphate Carboxylase Subunits (RBCS1-2B), an Hydroxypyruvate Reductase (HPR) and a Fumarase (FUM2) comprised the DEG subset from “carbon metabolism” sharing expression pattern with the glucosinolate pathway in coastal demes (Figure 7C). The last three KEGG pathways significantly enriched in the downregulated transcript set from *coreSAL* were “glyoxylate and decarboxylate metabolism” (19 DEGs), “glycine, serine and threonine metabolism” (10 DEGs) and “nitrogen metabolism” (8 DEGs) (Figure S6; Table S4). *PGLP1* (AT5G36700, from the glyoxylate decarboxylate metabolism) suffered x1.4 times more downregulation in coastal demes under *alkSAL*. PGLP1 is one of the core metabolite repair enzymes of plant photosynthetic carbon assimilation, as it degrades 2-phosphoglycolate (2PG) for its reconversion to Calvin cycle metabolites (Flügel et al., 2017). After 2PG degradation to glycolate, glyoxylate is generated, ultimately being catalyzed to glycine (Dellero et al., 2016). Thus, *coreSAL* inhibition of glyoxylate and decarboxylate transcripts in the studied demes is likely affecting glycine, serine, and threonine metabolism. In this regard, 10 DEGs enriched in the “glycine, serine and threonine metabolism” pathways were downregulated in the 4 demes under neutral and alkaline salinity (Table S6), and *AK3* (AT3G02020) was x1.6 times more downregulated in T6^CS^ and Ro2^CS^ when compared to V1^PC^ and LM2^PC^ under *alkSAL* (Figure S6). AK3 codes for an aspartate kinase that catalyzes the first step in aspartate-derived amino acids (Clark et al., 2015). *AK3* upregulation was also reported in A1 – carbonate tolerant deme – under bicarbonate exposure when compared to control, but not in T6^CS^ – carbonate sensitive deme (Pérez-Martín et al., 2021).

Finally, inhibition of transcripts from the α-CA and β-CA (Carbonic Anhydrases) (α-CA1, α-CA2, β-CA1, β-CA2 and β-CA4), and the glutamine synthetases 1;4 (GLN1;4) and 2 (GS) family proteins comprised the “Nitrogen metabolism” down-regulated set (Table S6). From these, *βCA-1* (At3G01500) and *α-CA1* (At3G52720) displayed x1.7 times more inhibition in T6^CS^ when compared to V1^PC^ and LM2^PC^ under *alkSAL* (Figure S6). βCA-1 is reported to regulate CO_2_ controlled stomatal movements in guard cells together with *βCA4* (AT1G70410). Previous electrophysiological assays reported *βCA4* reduction in T6^CS^ under bicarbonate conditions (Pérez-Martín, 2021), and *βca1ca4* mutant under NaHCO_3_ conditions show reduced photosynthetic efficiency (F_v_/F_m_) and increased total ion leakage (as reporter for cell death) under NaHCO_3_ treatment (Dąbrowska-Bronk et al., 2016). α-CA1 belongs to the α-CA family and is involved in the transformation of HCO^3−^ to CO_2_ in chloroplast stroma to supply it at the active site of Rubisco (Burén 2010).

Considering the high demand of Fe cofactors by photosynthetic enzymes (Kroh & Pilon, 2020), the above results raise the hypothesis that the defective Fe use efficiency in coastal demes under *alkSAL* (especially T6^CS^) leads to a stronger photosynthetic inhibition. This would result in the observed unbalance in carbon fixation and metabolism in T6^CS^ and Ro2^CS^, which ultimately would compromise their total energetic investment into secondary metabolism. In such a scenario, the stronger downregulation of PGLP1 (entry enzyme into one of the major pathways of primary plant metabolism) and the chloroplast carbonic anhydrases β-CA1 and α-CA1 in coastal demes under *alkSAL* is consistent with their reduced photosynthesis efficiency (F_v_/F_m_ values) and stronger downregulation of transcripts from photosynthesis (PSAH1-2), carbon fixation and metabolism (RBCS1-2B, HPR and FUM2). FUM2 (Fumarate hydratase 2 - AT5G50950, encoding a cytosolic chloroplast fumarase) would link primary metabolism to secondary metabolism (Figure S7), leading to the downregulation of all transcripts involved in glucosinolate metabolism and some transcripts from the glycine, serine and threonine metabolism reported to enhance alkalinity tolerance, like AK3. Likewise, the results confirm that a significant inhibition of glucosinolate signaling in coastal demes is caused in an alkalinity-specific manner, likely by decreased availability of ferrous ions (Fahey et al., 2001; Wittstock et al., 2002).

#### A reduced coreSAL DEG subset orchestrate contrasted plant response to neuSAL and alkSAL

All shared DEGs displaying opposite expression trends between neutral and alkaline salinity in more than 2 demes were analyzed to seek for contrasted transcriptomic responses among the studied demes potentially driving their differential performance under each salinity type (Figure 7D). *BGAL4* (AT5G56870), *DIN10* (AT5G20250) and *SEN1* (AT4G35770) exhibited contrasted expression trends among V1^PC^ and LM2^PC^ (down-regulated) and T6^CS^ and Ro2^CS^ (up-regulated) under *neuSAL* (Figure 8A). This suggests that activation of *BGAL4*, *DIN10* and/or *SEN1* is advantageous under *neuSAL* but not under *alkSAL*. Therefore, the genomic sequences of *BGAL4*, *DIN10* and *SEN1* in the 4 study demes were extracted from the Catalan demes germplasm Whole Genome Sequencing (WGS) data (Busoms et al., 2018) to identify sequence variation likely causal of the contrasted expression trends in the studied demes under *neuSAL* and *alkSAL* (Figure S8; Table S7). A SNP located only 4 bp upstream the 5’UTR at *BGAL4* promoter was consistent for all replicates of each deme and, moreover, differentiated coastal from non-coastal demes (Figure 8B).

**Figure 8.**
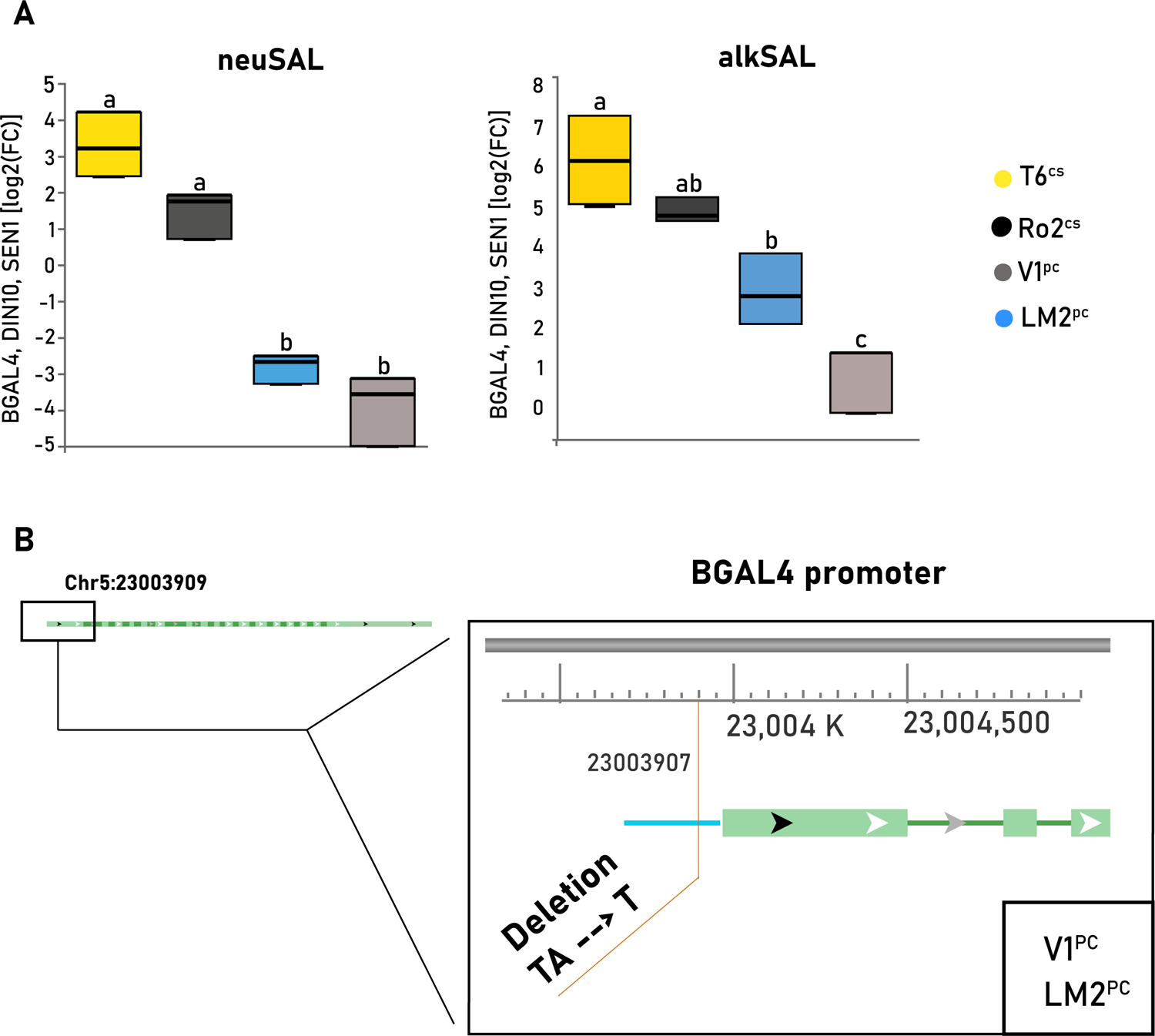
Characterization of *BGAL4* expression and sequence variation in *A. thaliana* demes. **(A)** Expression levels [mean ± SE log2(FC)] of *BGAL4, DIN10* and *SEN1* in each deme (n=3) and treatment. Letters indicate significant differences in mean expression values among demes (Tukey’s HSD, adj. *p*-value < 0.05). T6^CS^ (yellow), Ro2^CS^ (black), V1^PC^ (blue), and LM2^PC^ (grey). **(B)** SNP polymorphisms surrounding the *BGAL4* locus. Triangles: candidate SNPs position. Amino acid change is indicated below each SNP. Gene orientation is indicated with an arrow on the right. Exons are indicated with boxes and introns with lines connecting them. Chromosome positions are indicated at the top left (picture from https://www.arabidopsis.org/index.jsp). Reference and alternative allele nucleotide changes, variant type and deme bearing the SNP can be found in Figure S8.

BGAL4 encodes a glycosyl hydrolase regulated by SNF1-related Kinase 1 (SnRK1), a kinase involved in key plant stress responses (Peixoto et al., 2021). Degradation of structural polysaccharides in plant cell wall by cell wall-remodeling enzymes - like BGAL4 – forms part of the stress-regulated cell wall rearrangement (Moneo-Sánchez et al., 2016). A study comparing *A. thaliana* Wild Type vs salt-tolerant mutant roots exposed to NaCl stress reported higher expression of glycosyl hydrolases in the salt-tolerant mutant root, which was hypothesized to increase its potential of salt resistance (Guo et al., 2014). In contrast, β-galactosidases are reported to dramatically decrease enzymatic activity at high pH and to have pH optima ranging from 3.5 to 5.0 (Ross et al., 1994; Smith & Gross, 2000; Kotake et al., 2005, Hussien et al., 2021). Thus, enhanced upregulation of *BGAL4* in T6^CS^ and Ro2^CS^ could point to an enhanced activation of plant energy management functions under *neuSAL*, whereas a lack of ability to modulate BGAL4 activity for the prioritization of compounds more suitable under alkaline conditions could compromise their energy budget (Tsai & Schmidt, 2020). Moreover, the decline in photosynthesis and carbon fixation efficiency observed in coastal demes under *alkSAL* may cause further stimulation of the hydrolase enzymes, as these are triggered by sugar starvation (Lee et al., 2007). Transcription factor binding is the strongest contributor to variation in mRNA levels (Pai et al., 2015). This might be the case for *BGAL4*, where the SNP change in the promoter region of V1^PC^ and LM2^PC^ could contain cis-regulatory elements responsible for its consistently lower expression.

### 2. WGCNA Network

To combine transcriptomic and physiological datasets and detect possible correlations between genes determining traits of interest, all DEGs detected in each deme under *neuSAL* and *alkSAL* were retained for WGCNA unsigned co-expression network analysis. This involved a total of 11432 DEGs (11K WGCNA hereafter). The soft threshold power of 7 (β = 7) was selected according to the preconditions of approximate scale-free topology (Figure S9). WGCNA 11K analysis identified sixteen distinct co-expression modules which were assigned to different colors shown in the dendrogram (Figure S10). The obtained module-trait correlation matrix displayed two major contrasted patterns defined by correlation trends in 3 main traits: leaf Na^+^, leaf B and relative rosette biomass (Figure 9). This confirmed that correlation patterns observed in the performed multivariate analyses of physiological traits (PCA and MPC) were maintained when integrating co-expressed transcriptome data into trait correlations (WGCNA). Besides, most co-expression modules that negatively correlated with leaf Na^+^ but positively correlated with leaf B and sample biomass were also displaying positive correlation with leaf K^+^ and F_v_/F_m_ values, confirming their biological relevance in the context of salinity stress.

**Figure 9.**
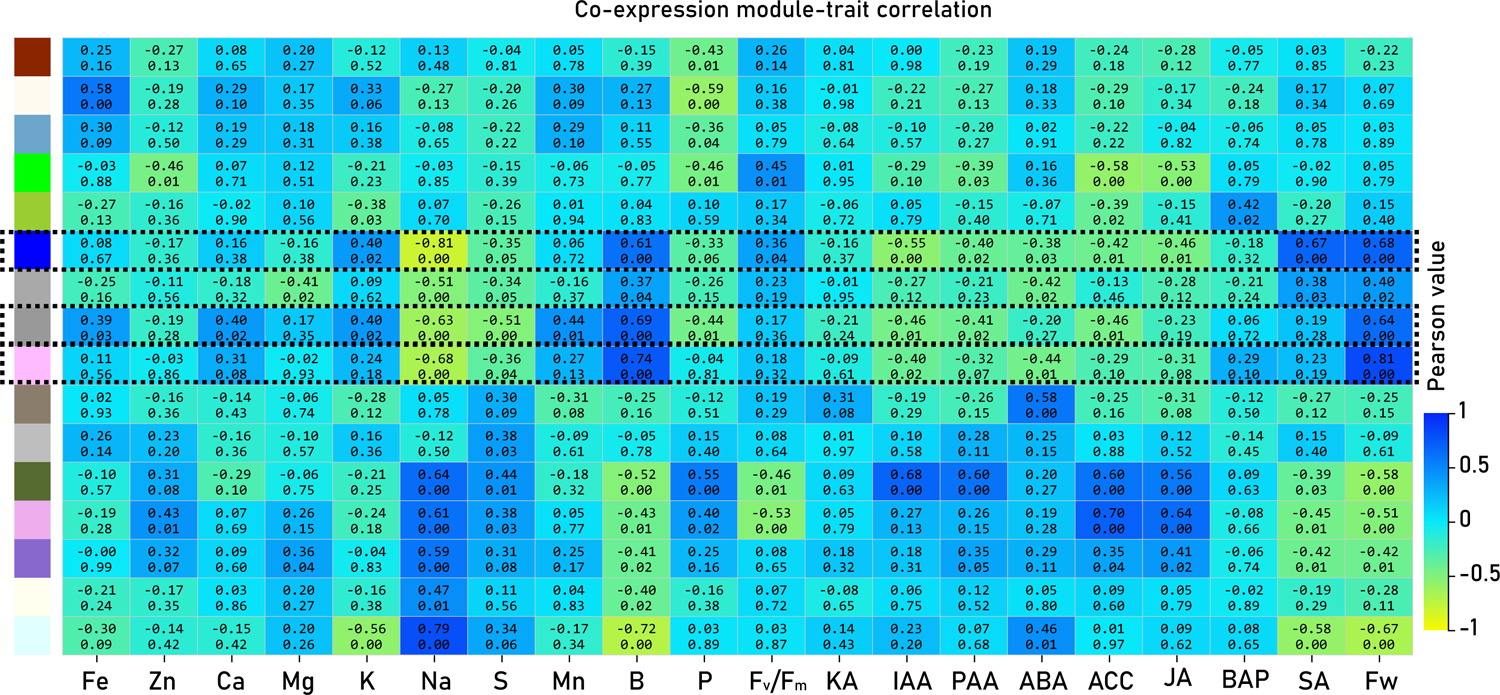
WGCNA analysis. Relationships of co-expression modules and analyzed physiological traits. Each row in the table corresponds to a co-expression module, and each column to a trait. Numbers in the matrix report the correlations of the corresponding module genes and each trait, with the p values below the correlations in parentheses. The table is color coded by correlation according to the color legend.

Considering all the above, modules showing significant negative correlation with Na^+^ and positive correlation with B, K^+^ or FW were picked for further analysis of their implication in tolerance mechanisms to salinity stress and GO and KEGG enrichment analysis of each selected co-expression module was performed for each significant selected module. After that, DEGs comprised in the significantly enriched pathways of each module were merged with the representative set of DEGs from *coreSAL*, *onlyNEUSAL* and *onlyALKSAL*, and results confirmed that DEGs selection criteria carried for transcriptome analysis – picking only DEGs detected in at least 3 out of the 4 demes comprised in the shared and the exclusive response to neutral and alkaline salinity - accurately reflected the behavior of the entire dataset (Table S8). Moreover, all the genes highlighted from previous analysis have also been identified in the 3 modules explored (Figure 10).

**Figure 10.**
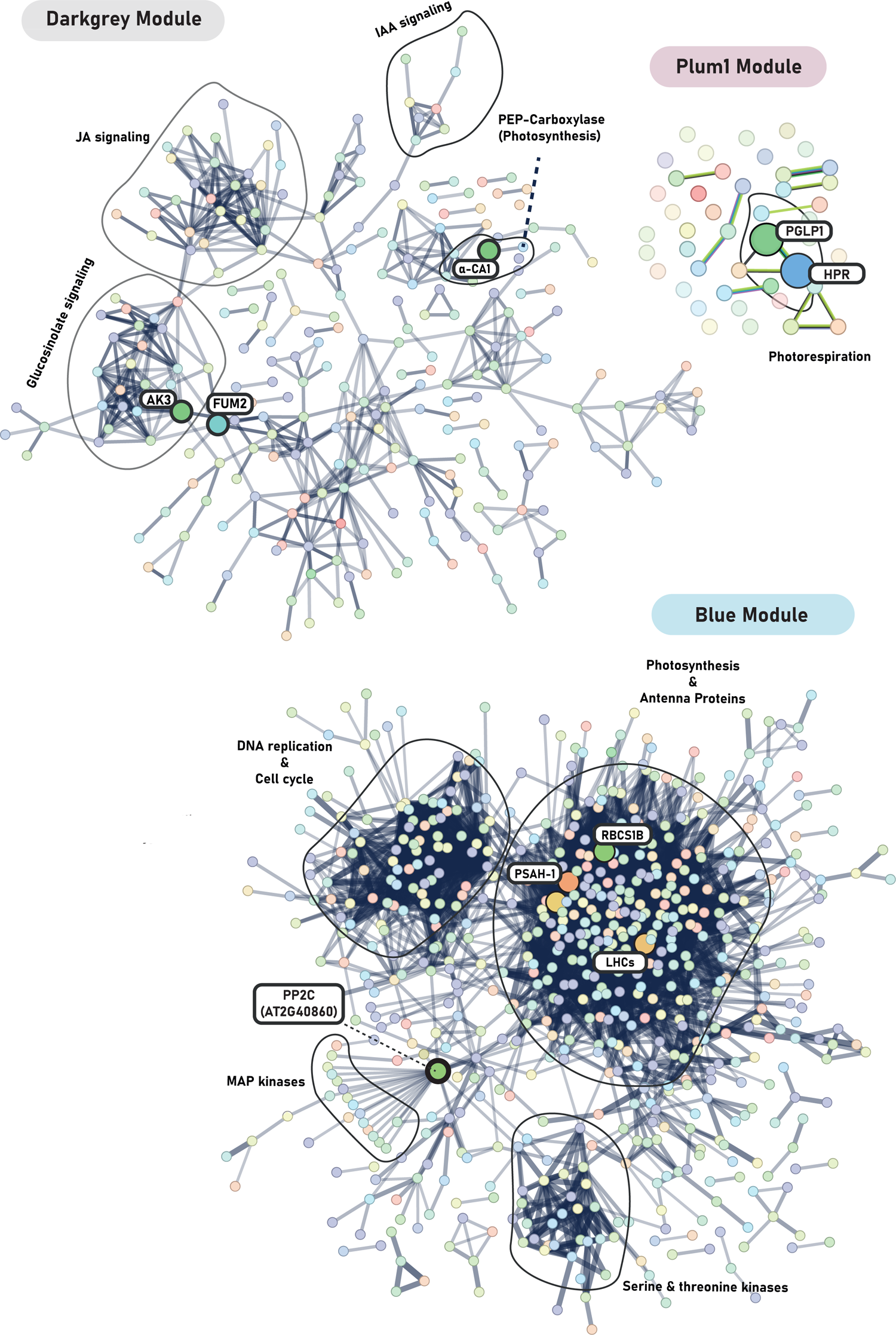
Overview of WGCNA co-expression modules significantly correlating to alkaline salinity tolerance traits. Protein-protein interaction (PPI) network of all DEGs comprised in darkgrey, plum1 and blue WGCNA co-expression modules, based on the STRING online database (https://string-db.org/). Proposed candidates identified in each module and appearing on each PPI Network are indicated, and main signaling pathways enriched per module encircled.

Overall, the combination of multivariate analysis on physiological traits, transcriptome analysis and WGCNA results prove that the selection criteria performed to assess effects of salinity in the studied demes are an accurate reflection of the whole dataset behavior. This combination provides knowledge on the hub biological processes and metabolic pathways that reflect the differential responses of neutral-salinity-adapted demes when compared to demes with tolerance to moderate alkaline-salinity. Moreover, it gives evidence supporting the involvement of key nutritional (K^+^ homeostasis, B sufficiency, Na^+^ uptake reduction and Zn limitation) and phytohormonal cues driving tolerance to salinity.

## Conclusions

By means of an integrative physiological, ionomic, metabolomic and transcriptomic approach, convergent and divergent biological pathways conferring shared and exclusive responses to neutral (*neuSAL*) and alkaline salinity (*alkSAL*) have been revealed in natural populations of *Arabidopsis thaliana*. From the analysis of up-regulated transcripts at the core of salinity, we conclude that activation of the protein synthetic machinery and changes in cell wall composition by FARs -mediated suberin deposition processes are common responses to salinity. Whereas under *neuSAL* antioxidant mechanisms are boosted mainly through the anthocyanin biosynthesis pathway, the phenylpropanoid and glutathione pathways are raised under *alkSAL*. Through screening of DEGs with contrasted expression profiles among demes, we propose that a smart regulation of BGAL4, a cell wall and tonoplast hydrolase, is key to tolerate different salt stresses, and a SNP detected in the 5’UTR of V1^PC^ and LM2^PC^ is proposed as a candidate for the differential regulation of gene expression levels. In contrast, dissection on the down-regulated transcripts at the *coreSAL* set confirms that tolerance to neutral salinity does not confer advantageous features under alkaline salinity. Key genes involved in the photorespiratory cycle – PGLP1, βCA1, RBCS1-2 -, organic acid accumulation – FUM2 -, and other secondary metabolism processes for abiotic stress resistance – AK3, glucosinolate biosynthesis genes – could be targets for alkaline salinity resistance breeding. WGCNA analysis supports that the presented candidates are comprised in co-expression modules significantly correlating with favorable responses to *alkSAL*. In this concern, increasing internal Fe use efficiency for maintenance of photosynthesis rate, and rising Ca, B and K accumulation for lowering Na-K, Na-Ca, Na-B ratios might be essential when facing alkaline salinity stress. Moreover, the enhanced activation of ABA-dependent signaling is highlighted as a key trait for plant performance under *alkSAL*, likely by PYL/PYRs binding to PP2Cs and consequent activation of H^+^ATP-ase activity, allowing apoplastic acidification. In turn, sustained ABA accumulation and signaling through WRKYs regulation are shown exclusively in coastal demes T6^CS^ and Ro2^CS^ under *neuSAL* and are proposed to be adaptive strategies for these *A.thaliana* demes.

## Material & Methods

### Plant material

Seeds of 2 coastal-siliceous (T6^CS^ and Ro2^CS^) and 2 intermediate-calcareous (V1^PC^, LM2^PC^) *A. thaliana* demes were collected from the natural habitat (Figure S1) and propagated in the laboratory. The flowchart in Figure S2 summarizes the experimental procedures of the study.

### Irrigation experiment

Seeds were surface sterilized by soaking them in 30% (v/v) commercial Clorox bleach and 1 drop of Tween-20 for 15 min and rinsed 5 times in 18 MΩmilli-Q sterile water. Seeds were stratified at 4°C for 1 week in the dark and sowed on wet soil. Pots were covered with PVC film until the seedlings germinated. Pots with germinated seedlings were placed in a growth chamber (Conviron CMP5090) with 8 h light / 16 h dark photoperiod, an irradiance of 80 mmol·m^-2^·s^-1^ and a constant temperature of 22 °C. Plants were watered with 0,5-strength Hoagland solution every 3 days. After 2 weeks, all plants were irrigated every 3 days with 0,5-strength Hoagland solution containing 0,5-strength Hoagland solution alone (control), with 100 mM NaCl (neutral salinity: ***neuSAL***) or with 85 mM NaCl and 15 mM NaHCO_3_ at pH 8.3 (alkaline salinity, ***alkSAL***) for two weeks. The Rosette Diameter (RD) and the PSII efficiency of each sample were measured after two weeks of treatment. After harvest, samples for ionomic analysis were dried and samples for RNA extraction and hormone determination were immediately frozen in liquid nitrogen and stored at -80 °C.

### Leaf nutrition

Plant tissues were sampled by removing 2–3 leaves or cutting roots (1–5 mg dry weight). They were then washed with 18 MΩ water before placing them in Pyrex digestion tubes. The sampled plant material was dried for 2 days at 60 °C and weighed before open-air digestion in Pyrex tubes using 0.7 mL concentrated HNO_3_ at 110 °C for 5 h in a hot-block digestion system (SC154-54-Well Hot Block, Environmental Express, Charleston, SC, United States). Concentrations of the selected elements (Ca, K, Mg, Na, P, S, Mo, Cu, Fe, Mn, Zn) were determined by ICP-MS (Perkin ElmerInc., ELAN 6000, MA, United States) or ICP-OES (Thermo Jarrell-Ash, model 61E Polyscan, England) (n=6 per deme and treatment).

### PSII efficiency

The maximal photochemical efficiency of PSII was calculated from the ratio of variable (Fv) to maximum (Fm) fluorescence [F_v_/F_m_ = (Fm - F_o_)/F_m_] using a MINI-PAM-II/B portable chlorophyll fluorometer (Walz) connected to an Arabidopsis Leaf Clip holder (model 2060-B). The minimum fluorescence yield (F_o_) was measured under measuring light (650 nm) with very low intensity (0.8 mmol m^-2^ s^-1^). To estimate the maximum fluorescence yield (F_m_), a saturating pulse of white light (3000 mmol m^-2^ s^-1^ for 1 s) was applied. Leaves were dark-adapted for 30 min before measurements and the analysis was performed in a dark room with stable ambient conditions (n=6 per deme and treatment).

### Endogenous phytohormones

Phytohormones extraction was done according to the protocol of Pan et al. (2008) with modifications from Delatorre et al. (2017). The phytohormones analyzed were abscisic acid (ABA), indoleacetic acid (IAA), 1-aminocyclopropane-1-carboxylic acid (ACC), jasmonic acid (JA), and salicylic acid (SA). The following deuterated standards were employed: ABA-^2^H6, IAA-d5, ACC-d4, JA-d5, and SA-d6. All of them were supplied by Olchemim Ltd. (Czech Republic) except SA-d6 obtained from Sigma Aldrich (St. Louis, MO, USA). Briefly, 0.05 g fresh weight of leaf samples were homogenized in liquid N_2_ and extracted with 250 µL of extraction buffer (2-propanol/Mili Q water/Hydrochloric acid; 2:1:0.002; v/v/v) in 1.5 mL Eppendorf tubes (Eppendorf Safe Lock England) and agitated by repeated inversion (60 rpm) for 20 min, at 4 °C, in the dark. The resulting homogenate was transferred to a Teflon tube (Thermo Scientific, England) with 450 µL dichloromethane and reextracted by repeated and inverted agitation for 30 min, at 4 °C in the dark. Three phases were obtained, an aqueous, a material debris and an organic phase. The organic phase (bottom) was recovered, and the intermediate debris phase was also re-extracted. The combined extracts were evaporated under N_2_ flow to remove propanol and resuspended 150 µL of 100% methanol. Resuspended extracts were filtered through a 0.22 µm cellulose acetate Spin-X centrifuge tube filter (Costar, Corning Inc., Salt Lake City, USA). Samples were injected into a QTrap4000 LC–ESI(−)–MS/MS system following the method described by Segarra et al. (2006) and modified by Llugany et al. (2013) using HPLC Agilent 1100 (Waldrom, Germany) on a Discovery C18 2.1 × 150 mm ID, 5 μm column (Supelco, Bellefonte, USA) at 50 °C, a constant flow rate of 0.6 mL min−1 and 10 μL injected volume. MS/MS experiments were performed on an API 3000 triple quadrupole mass spectrometer (PE Sciex, Concord, Ontario, Canada). All the analyses were performed using the Turbo Ionspray source in negative ion mode for SA, JA, ABA, IAA, and PAA and in positive mode for ACC. Quantification was performed by injection of extracted and spiked samples in multiple reaction monitoring (MRM) mode (n=6 per deme and treatment).

### Transcriptome profiling

#### Total RNA Exaction and RNA-Seq Library Preparation

Total RNA was extracted using Maxwell plant RNA kit (Promega Corporation, Madison, WI, USA) following the manufacturer’s instruction. The RNA concentration was measured using Qubit® 2.0 Fluorometer (Invitrogen). 4 μg total RNA was used to prepare RNA-seq libraries using the TruSeq Stranded mRNA Library Prep Kit (Illumina, San Diego, US). Strand specific paired-end mRNA sequencing was performed on DNBSEQ platform at the Next-Generation Sequencing Core of the BGI Genomics Service (Hong Kong, China) (n=3 per deme and treatment).

#### RNA-Seq analysis

Bowtie2 (Langmead & Salzberg, 2012) was used to map the clean reads to the reference gene sequence (transcriptome), and then used RSEM (Li & Dewey, 2011) to calculate the transcript abundance of each sample. Quality check of raw sequence data was performed using Fastqc. Reads containing adapters and poly-N and low-quality reads from raw data were removed with Qualimap. Simultaneously, Q20 (percentage of bases with the quality score greater than 20; sequencing error rate less than 1%), Q30 (percentage of bases with the quality score greater than 30; sequencing error rate less than 0.1%), and GC (or guanine-cytosine) content of the clean data were calculated. Paired-end reads were mapped to the TAIR10 reference genome using the STAR aligner and the .sam output of STAR was then converted to its compressed format .bam and sorted by gene identifier with samtools.

The overlap of reads with annotation features found in the reference .gtf was calculated using HT-seq (Putri et al., 2022). The output computed for each sample (raw read counts) were imported into BGI data visualization system (https://www.bgi.com/global/service/dr-tom) and parsed using DESeq2 (v 1.8.1). Raw counts were normalized using DESeq2’s function “rlog,” which normalises sequences according to library size to make them comparable among replicates and transforms the original count data to the log scale. The DEseq2 method is based on the principle of negative binomial distribution (Love et al., 2014). This method was used to perform differential gene expression analysis (DEG) with an absolute |FC| > 2. The *p*-values were corrected for multiple error testing according to Benjamini-Hochberg (*q*-value) and *q*-value threshold of 0.05 was set. According to the results of differential gene detection, the R package *pheatmap* was used to perform hierarchical clustering analysis and extract expression trends on the union set differential genes.

#### Gene functional annotation

Gene Onthology (GO) functional significant enrichment analysis - used to analyze molecular functions, cellular components and biological processes - was performed by mapping all candidate genes to each entry in the Gene Ontology database (http://www.geneontology.org/). A hypergeometric test was applied to find the GO function significantly enriched in candidate genes compared to all background genes of the species. *p*-value was calculated using the basis function phyper (https://stat.ethz.ch/R-manual/R-devel/library/stats/html/Hypergeometric.html) of R and then corrected for multiple error testing according to Benjamini-Hochberg (https://bioconductor.org/packages/release/bioc/html/qvalue.html). Finally, a *q*-value threshold of 0.05 was set, and the GO term that satisfied this condition was defined as the GO term that was significantly enriched in candidate genes. Kyoto Encyclopedia of Genes and Genomes (KEGG) pathway-based enrichment analysis – which links gene sets with a network of interacting molecules in the cell, such a pathway or complex - was performed following the same methodology and significance threshold than for GO functional enrichment analysis. Pathways significantly enriched in differentially expressed genes were identified and visualized through BGI data visualization system (https://www.bgi.com/global/service/dr-tom)

#### Protein-Protein Interaction (PPI) Network Analysis

Protein–protein Interaction (PPI) networks of differential expressed genes were performed on STRING version 11.0. Active interaction sources, including text mining, experiments, databases, and co-expression as well as species limited to “*Arabidopsis thaliana*” and an interaction score > 0.4 were applied to construct the PPI networks.

#### WGCNA network analysis

Weighted gene co-expression network analysis (WGCNA) was conducted with the “WGCNA” package (Langfelder and Horvath, 2008) on BGI’s Dr.Tom system (http://biosys.bgi.com). Gene expression file and trait file were combined, and the soft thresholding power (β value) was filtered based on the calculation of scale-free topological fit index and mean connectivity. The topological overlap matrix (TOM) was constructed based on the topological overlap between pairwise genes, and hierarchical clustering analysis was performed. Cluster dendrogram plot and clustering tree of co-expression modules of DEGs were created. The co-expression relationships among different modules were analyzed and modules with high similarity were merged at a similarity threshold of 0.25 and a minimum module size of 20 genes.

#### Statistical Analysis

All statistical analyses were performed using JMP SAS software (SAS Institute, Cary, NC, United States). Significant differences for multiple comparisons were determined by one-way or two-way ANOVA as indicated in figure legends, treatment or treatment relative to control (neuSAL/Control; alkSAL/Control) and deme (T6^CS^, Ro2^CS^, V1^PC^, LM2^PC^) or deme origin [coastal-siliceous_(T6+Ro2)_; intermediate-calcareous_(V1+LM2)_] as independent factors. Comparisons among demes were followed by post hoc Tukey HSD test (adjusted *p*-value < 0.05). Multivariate Correlation Analysis (MCA) was performed to determine the level of association among the measured physiological traits for each deme and treatment. Pearson’s correlation coefficients were calculated for all pairs of measured parameters under neuSAL and alkSAL at a level of 5% significance. Principal Component Analysis, eigenvalues and relative proportion of the variance explained by each trait included in MCA were calculated.

## Acknowledgements

Special thanks to Rosa Padilla for processing ICP and HPLC samples. This research was supported by the Spanish Ministry of Science and Innovation (MICINN) (PID 2019 104000 RB I00).

## Authors contribution

MJ-A performed the lab experiments. MJ-A, LP-M, GE-O and AL-V performed data analysis. MJ-A, SB, LP-M, AG, ML and CP conceived the study. All authors contributed to the writing of the manuscript.

---

**Figure S1.**
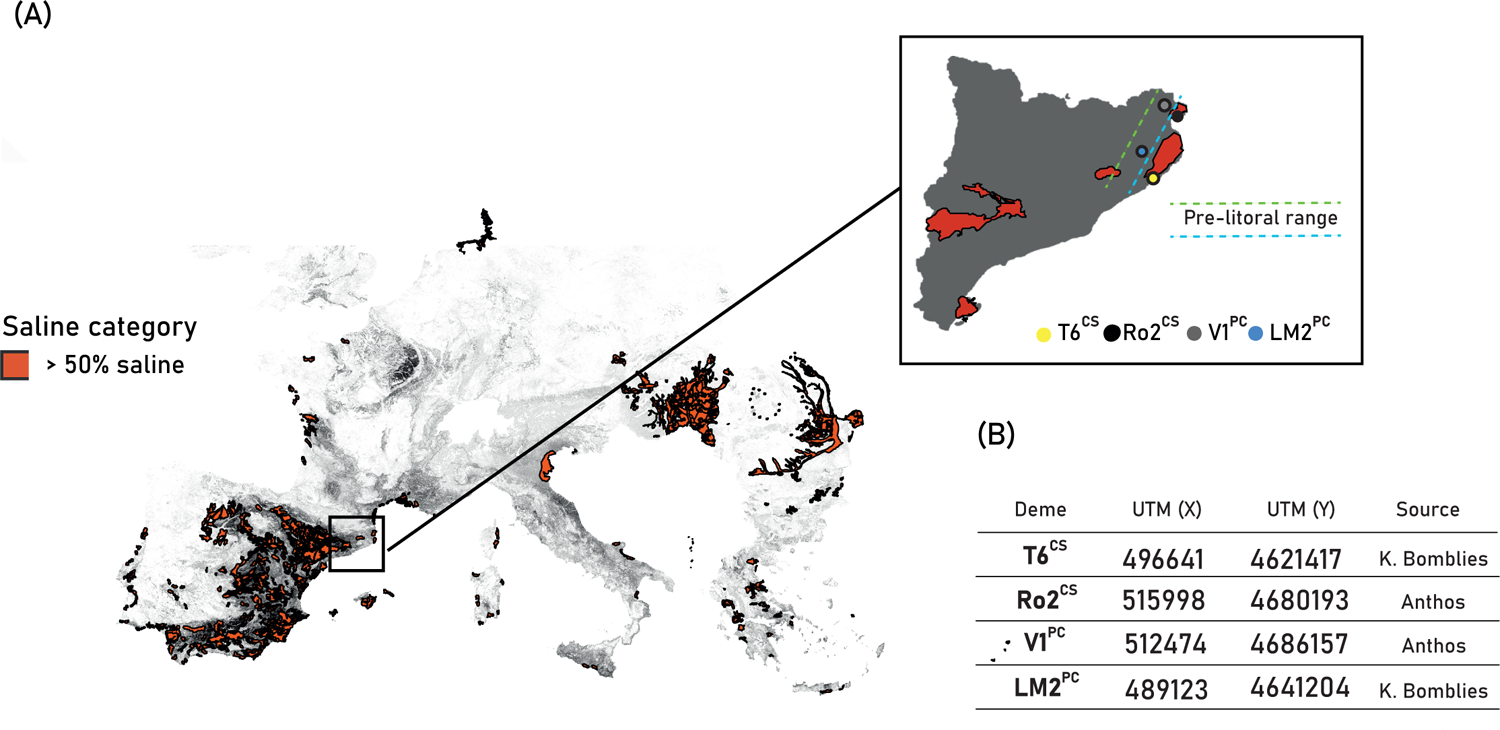
Plant material and soil of origin. **(A)** Distribution of saline soils in Europe. Red area is categorized as saline (>50% surface). Frame shows studied demes in Catalonia’s map **(B)** Table lists deme UTM coordinates and source.

**Figure S2.**
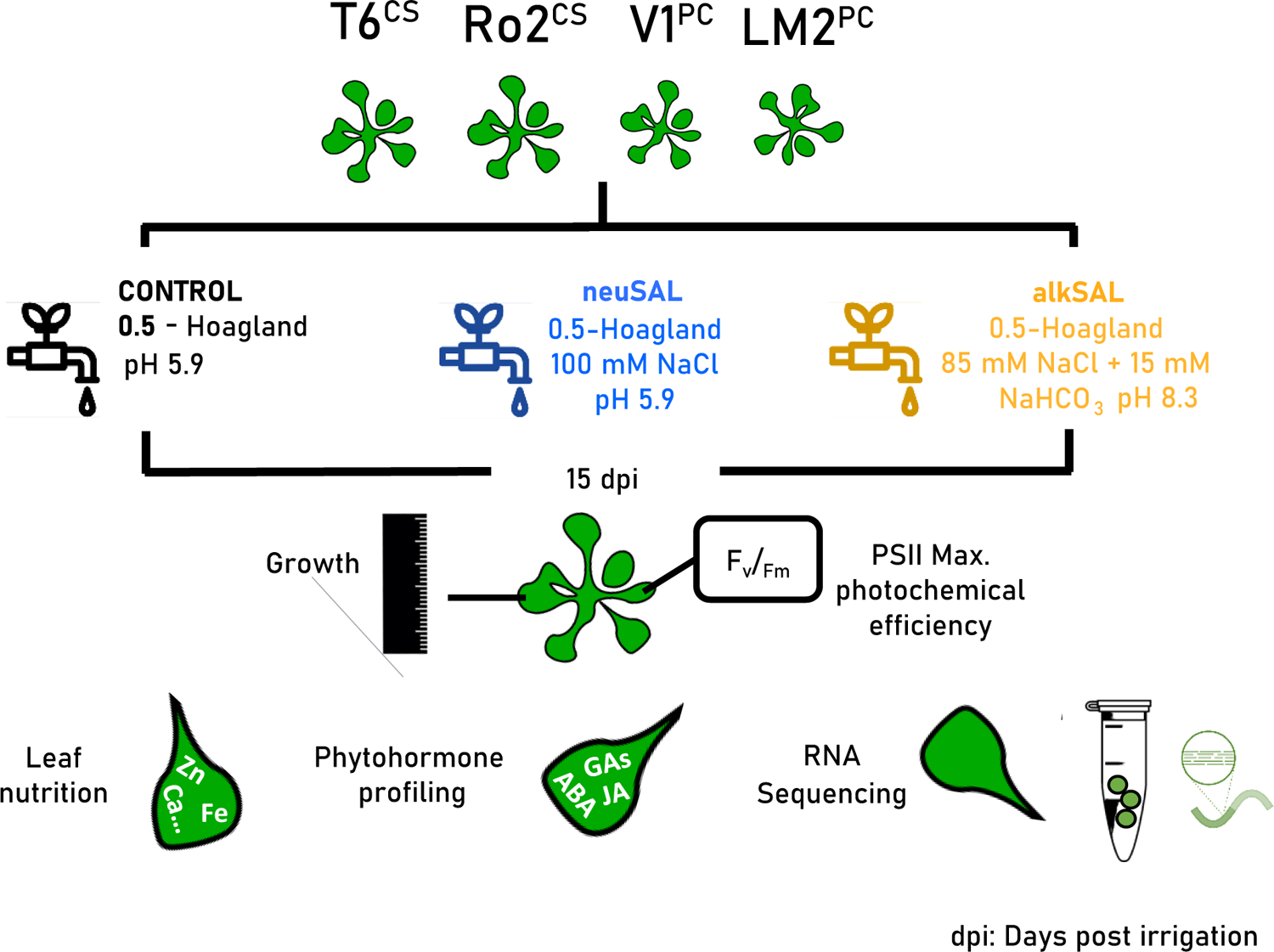
Workflow. Schematic overview of the compared sample growth conditions and performed analyses.

**Figure S3.**
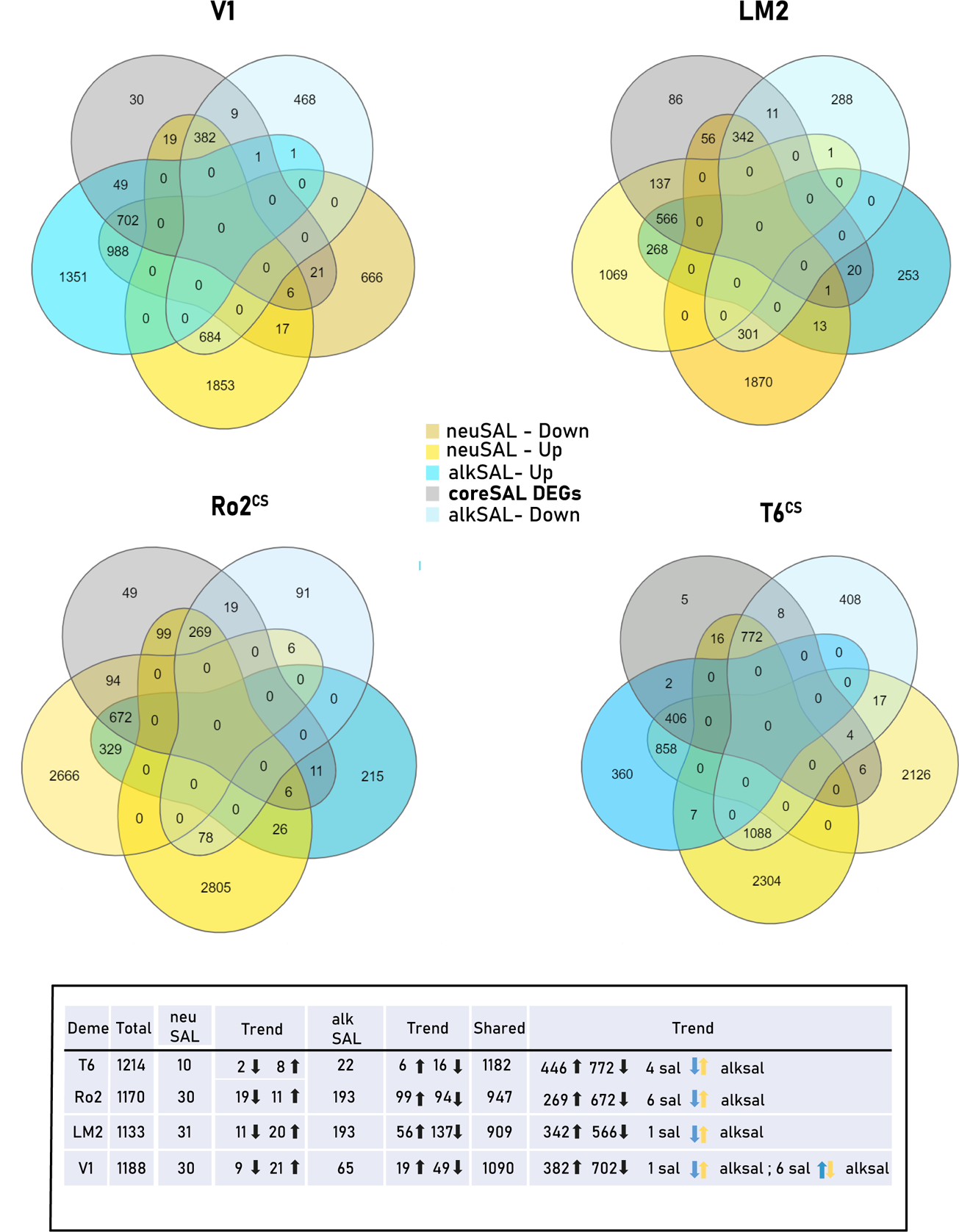
Contribution of each deme to the coreSAL DEGs set. Venn diagrams show overlapping DEGs in *coreSAL* (grey) and total DEGs per deme (yellow: total DEGs from *neuSAL* treatment; blue: total DEGs from *alkSAL* treatment). Table summarizes how many coreSAL DEGs are altered in each deme (“Total”), the salinity type causing the alteration (“*neuSAL*”, “*alkSAL*”, “*Shared*”) and number of upregulated, downregulated and mixed-trends DEGs.

**Figure S4.**
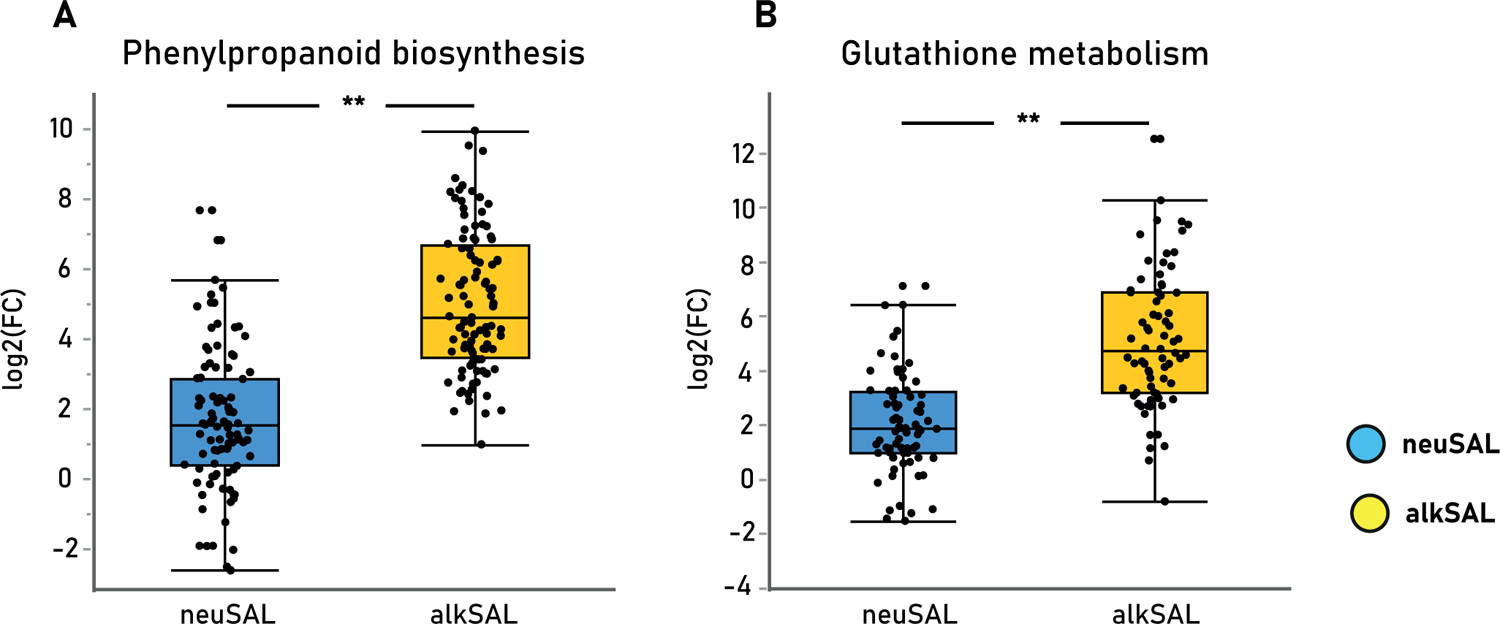
Dissection on significantly enriched pathways from *onlyALKSAL* upregulated DEGs. Expression levels [mean ± SE log2(FC)] of DEGs from **(A)** “Phenylpropanoid biosynthesis” (n=100) and **(B)** “Glutathione metabolism” (n=72) pathways in all demes among salinity treatments. Asterisks indicate significant differences in mean expression values (Student *t*-test, adj. *p*-value < 0.05; *: adj. *p*-value < 0.05; **: adj. *p*-value < 0.01) among treatments: ***neuSAL*** (neutral salinity in blue), ***alkSAL*** (alkaline salinity in yellow).

**Figure S5.**
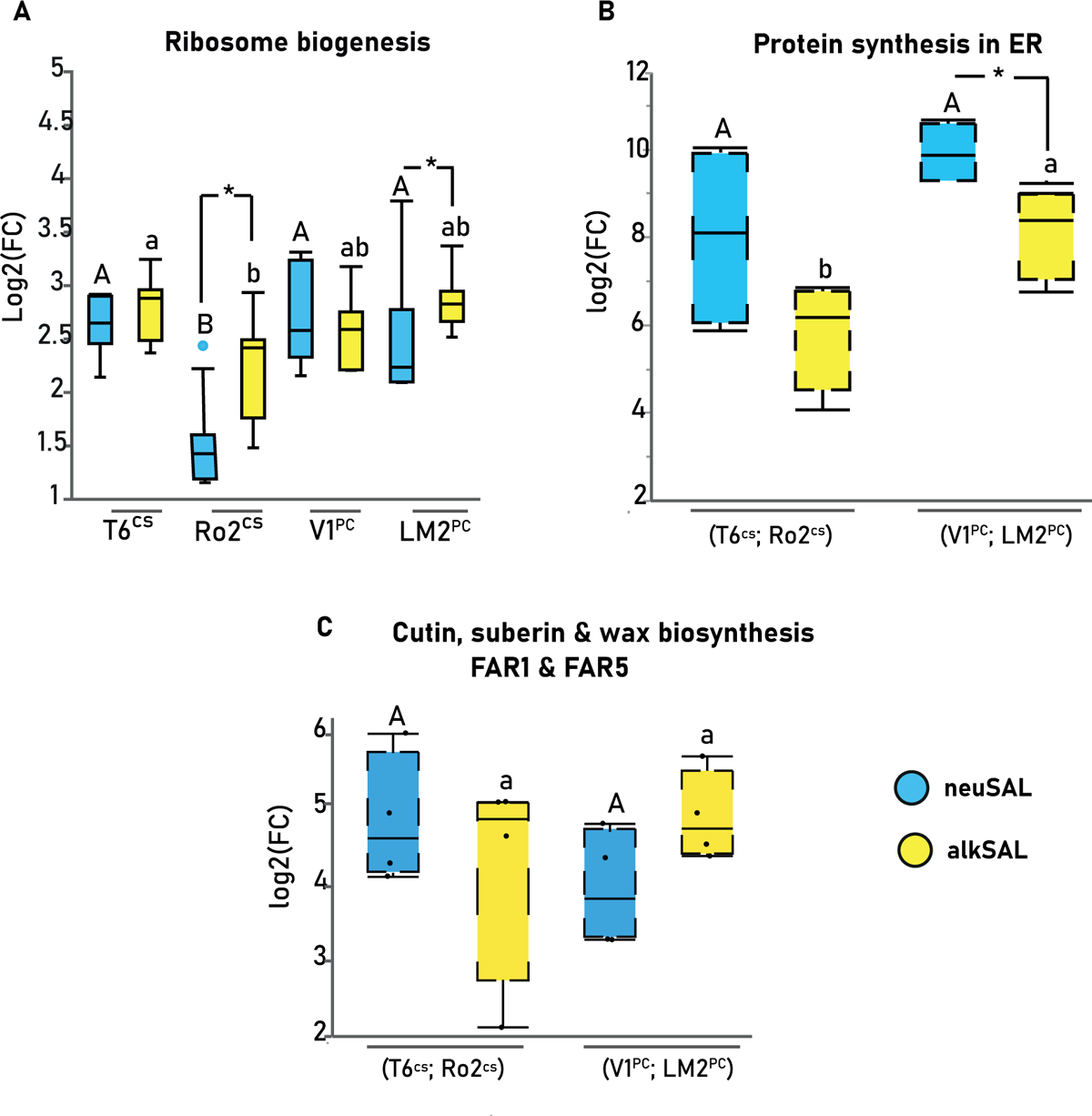
Dissection on significantly enriched pathways from *coreSAL* upregulated DEGs under neutral salinity (*neuSAL* in blue) and alkaline salinity (*alkSAL* in yellow) treatments. **(A)** Expression levels [mean ± SE log2(FC)] of DEGs from “Ribosome biogenesis” pathway in each deme and treatment (n=7). Letters indicate significant differences in mean expression values among demes under *neuSAL* (capital) and *alkSAL* (lower case) (Tukey’s HSD, adj. *p*-value < 0.05). Asterisks indicate significant differences in mean expression values among treatments for each deme (Student *t*-test, adj. *p*-value < 0.05 **(B)** Expression levels [mean ± SE log2(FC)] of DEGs from “Protein synthesis in ER” pathway in each location group (T6^CS^;Ro2^CS^/V1^PC^;LM2^PC^) and treatment (n=4). Letters indicate significant differences in mean expression values among location groups under *neuSAL* (capital) and *alkSAL* (lower case) (Student *t*-test, adj. *p*-value < 0.05). Asterisks indicate significant differences in mean expression values among treatments for each location group (Student *t*-test, adj. *p*-value < 0.05; *: adj. *p*-value < 0.05; **: adj. *p*-value < 0.01). **(C)** Expression levels [mean ± SE log2(FC)] of DEGs subset (FAR1 and FAR5) from “Cutin, suberin & wax biosynthesis” pathway in each location group (T6^CS^;Ro2^CS^/V1^PC^;LM2^PC^) and treatment (n=4). Letters indicate significant differences in mean expression values among location groups under *neuSAL* (capital) and *alkSAL* (lower case) (Student *t*-test, adj. *p*-value < 0.05).

**Figure S6.**
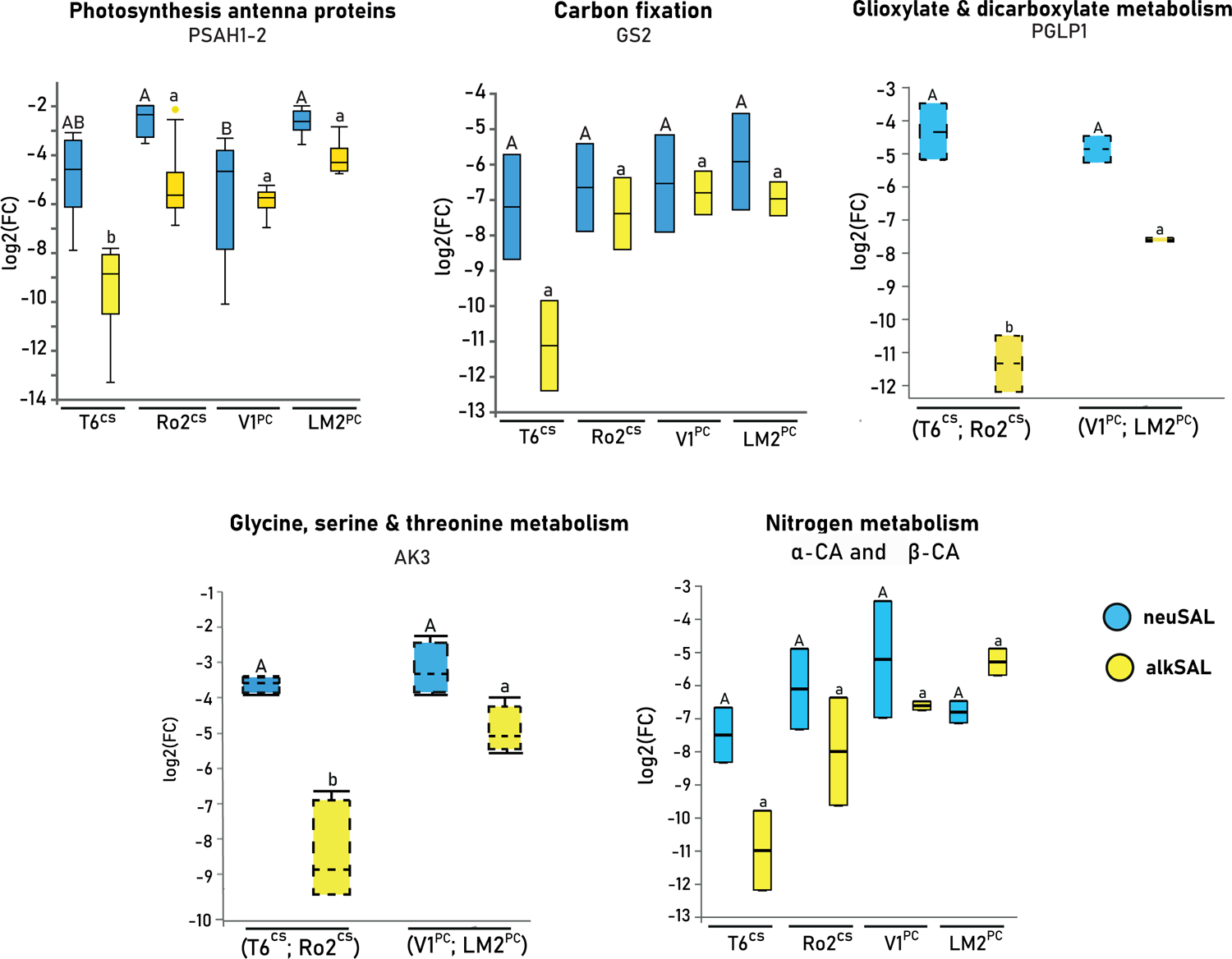
Dissection on significantly enriched pathways from *coreSAL* downregulated DEGs under neutral salinity (*neuSAL* in blue) and alkaline salinity (*alkSAL* in yellow**) treatments.** Expression levels [mean ± SE log2(FC)] of DEGs from **(A)** “Photosynthesis” (n=2), **(B)** “Photosynthesis Antenna proteins” (n=6), **(C)** “Glucosinolate biosynthesis” (n=9), **(D)** “Carbon metabolism” (n=3), **(E)** “Carbon fixation” (n=2), **(F)** “Glioxylate and dicarboxylate metabolism”, **(G)** “Glycine, serine and threonine metabolism” and **(H)** “Nitrogen metabolism” pathways in each deme and treatment. Letters indicate significant differences in mean expression values among demes under *neuSAL* (capital) and *alkSAL* (lower case) (Tukey’s HSD, adj. *p*-value < 0.05).

**Figure S7.**
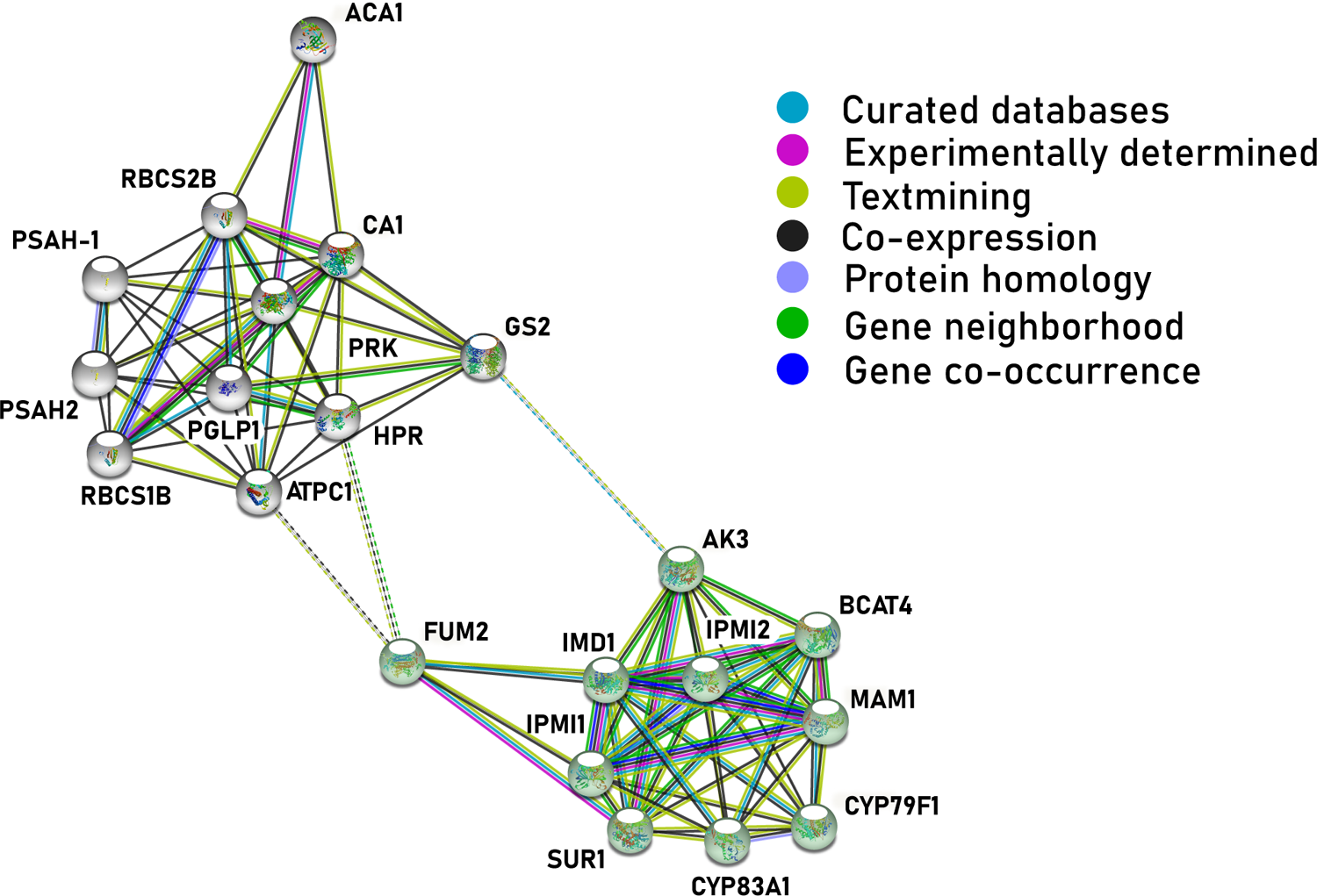
Key DEGs in the inhibition response of the studied demes. Protein-protein interaction (PPI) network of the down-regulated DEGs from enriched KEGG pathways in the *coreSAL* response. Performed at the STRING online database (https://string-db.org/).

**Figure S8.**
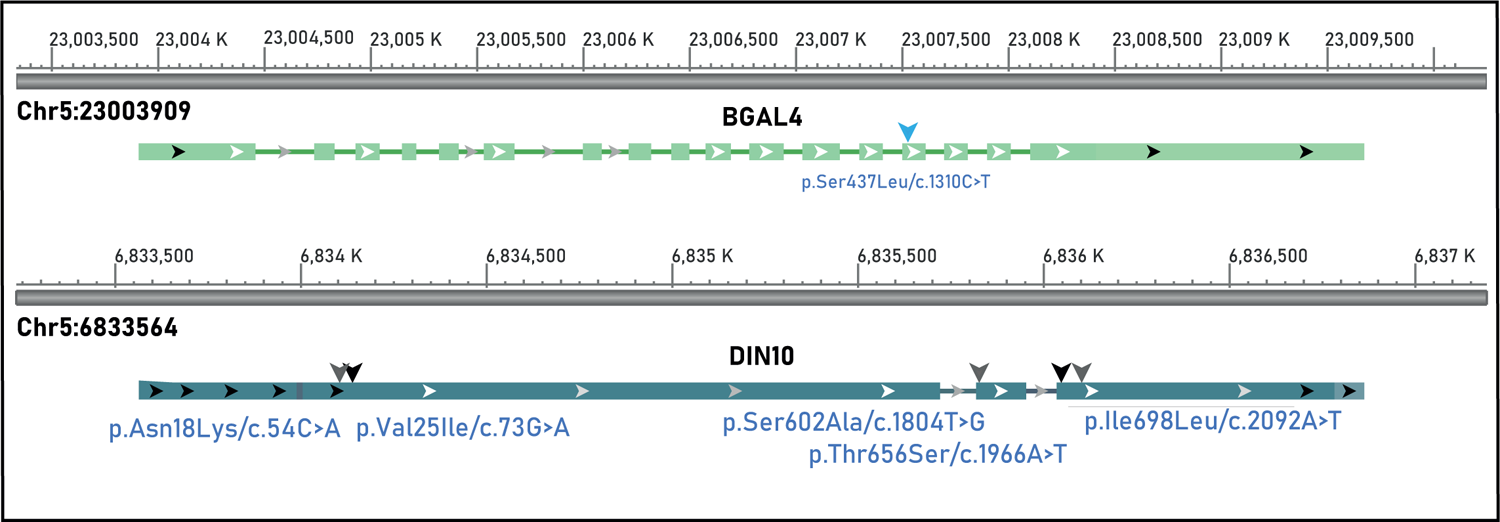
Gene models for BGAL4 and DIN10. Location of the SNPs identified at the coding sequence.

**Figure S9.**
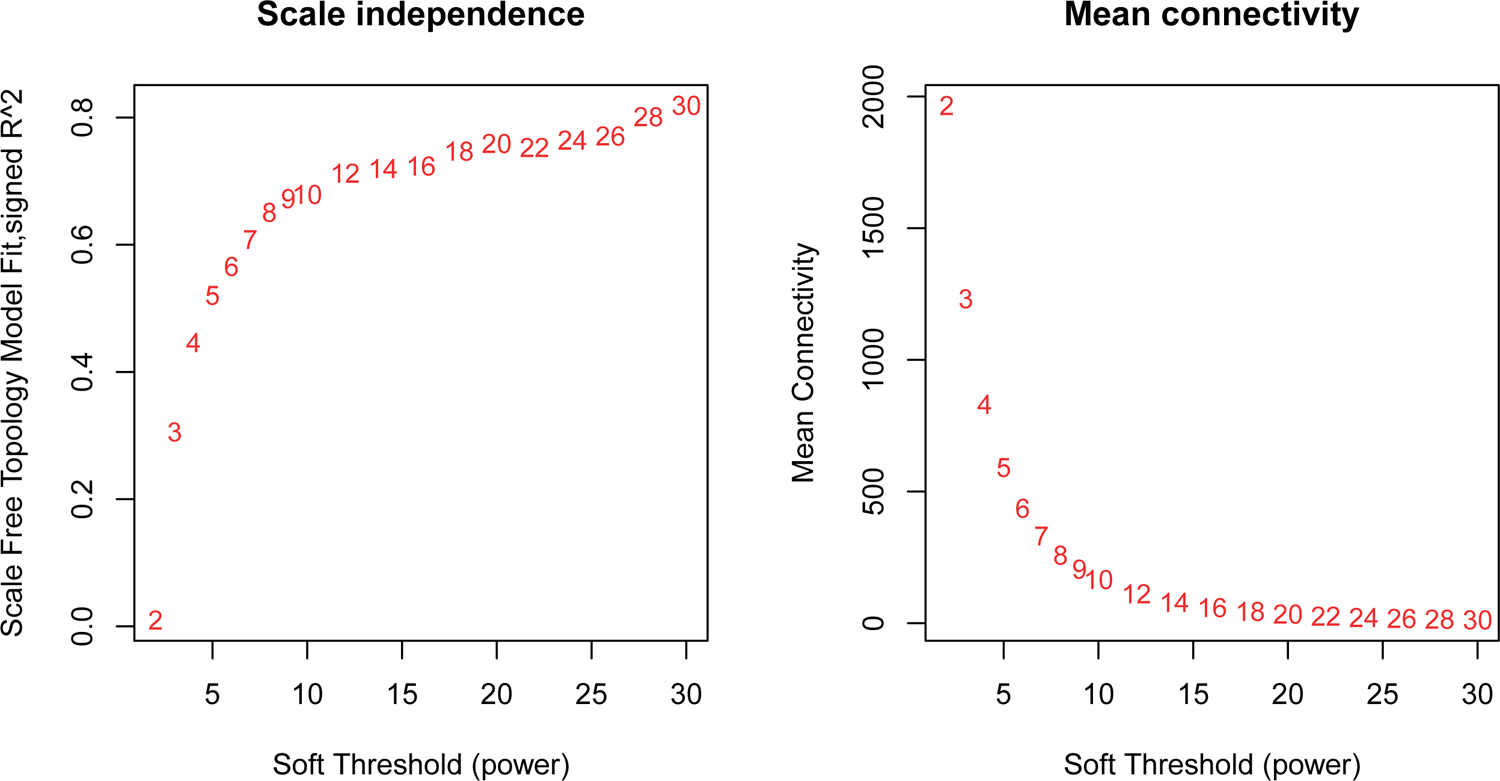
WGCNA analysis. Soft thresholding Network topology for different soft-thresholding powers. Numbers in the plots indicate the corresponding soft thresholding powers.

**Figure S10.**
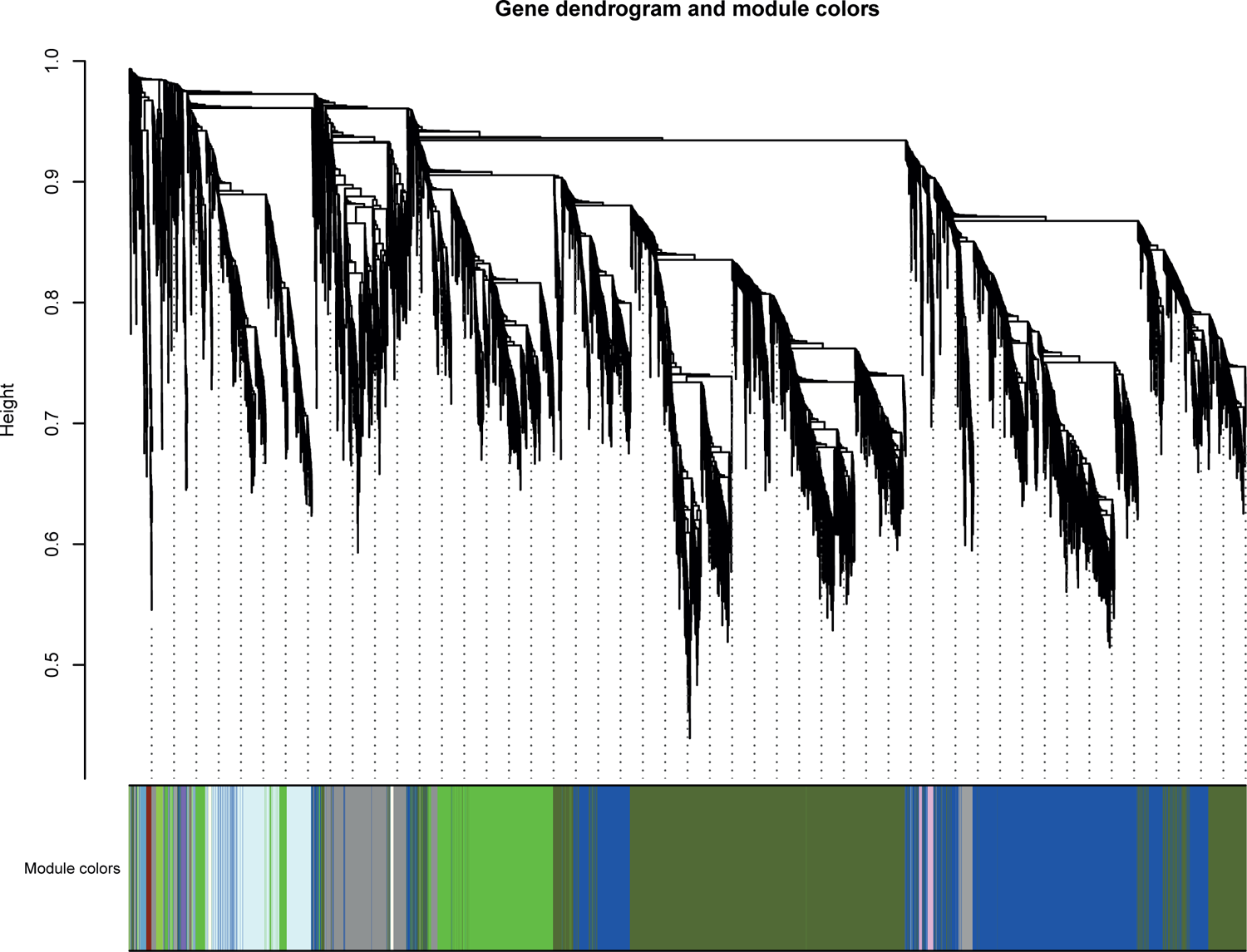
WGCNA analysis. Gene modules identified by Weighted gene co-expression network analysis. Gene dendrogram obtained by clustering the dissimilarity based on consensus topological overlap with the corresponding module colors indicated by the color row. Each colored row represents a color-coded module which contains a group of highly connected genes.

**Figure S11.**
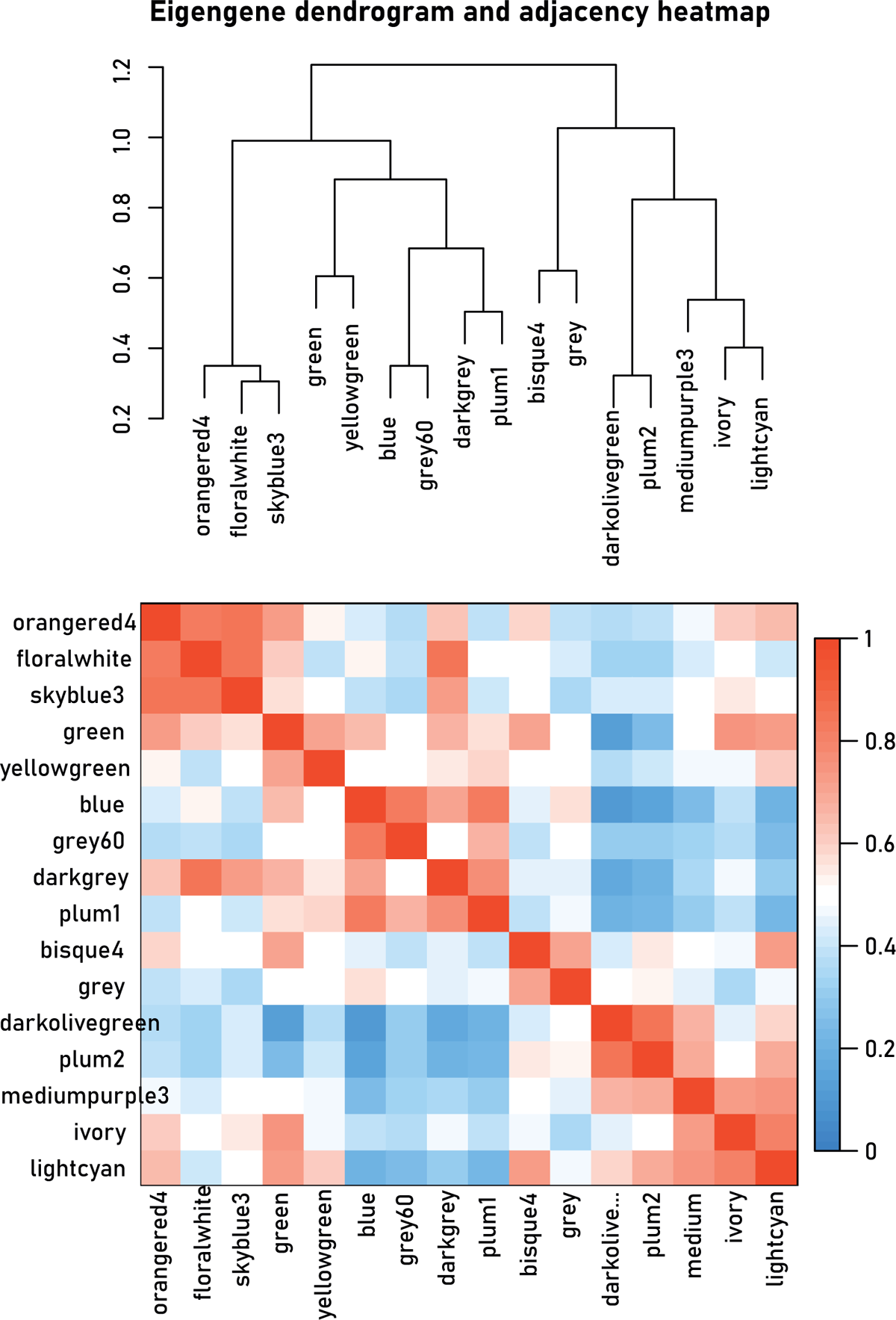
WGCNA analysis. Eigengene dendrogram and adjacency heatmap. Dendrogram of consensus module eigengenes obtained by WGCNA on the consensus correlation. The red line is the merging threshold, and groups of eigengenes below the threshold represent modules whose expressions profiles should be merged due to their similarity. Heatmap plotting adjacencies of modules. Red represents high adjacency (positive correlation) and blue represents low adjacency (negative correlation).

**Figure S12.**
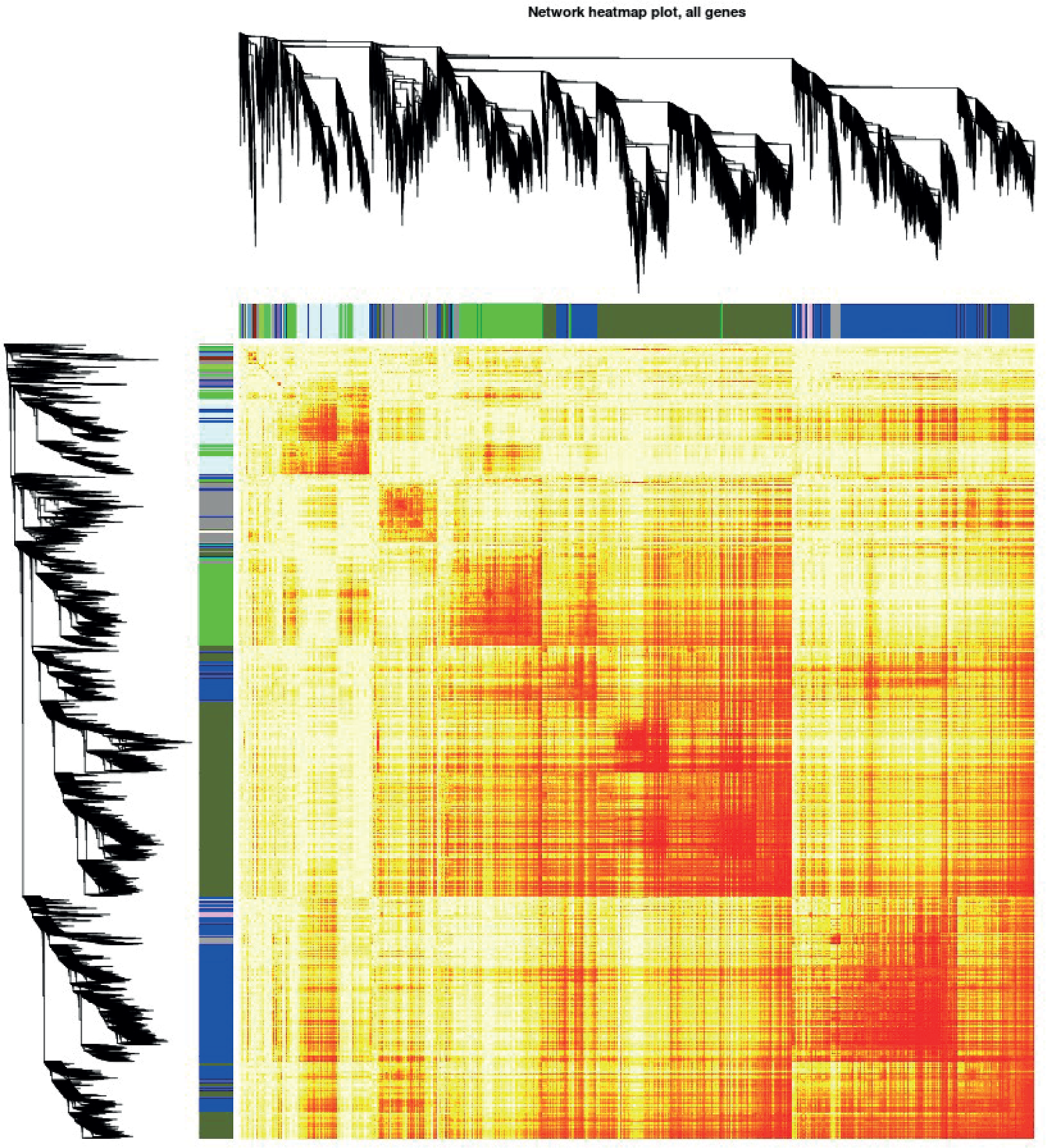
WGCNA analysis. Topological Overlap Matrix (TOM) showing the interaction of co-expression genes and the cluster dendrogram of 1,000 randomly selected genes. The intensity of the yellow inside the heatmap represents the degree of overlap (orange: high; yellow: low).

